# The RNA-binding protein SFPQ preserves long-intron splicing and regulates circRNA biogenesis

**DOI:** 10.1101/2020.09.04.280750

**Authors:** Lotte Victoria Winther Stagsted, Eoghan Thomas O’Leary, Thomas Birkballe Hansen

## Abstract

Circular RNAs (circRNAs) represent an abundant and conserved entity of non-coding RNAs, however the principles of biogenesis are currently not fully understood. To elucidate features important for circRNA production, we performed global analyses of RNA-binding proteins associating with the flanking introns of circRNAs, and we identified two factors, SFPQ and NONO, to be highly enriched with circRNAs. Using transient knockdown of both proteins in two human cell lines followed by total RNAseq, we found a subclass of circRNAs with distal inverted *Alu* elements and long introns to be highly deregulated upon SFPQ knockdown. In addition, SFPQ depletion leads to increased intron retention with concomitant induction of cryptic splicing prevalent for long introns causing in some cases premature transcription termination and polyadenylation. While SFPQ depletion has an overall negative effect on circRNA production, premature termination is not the main causative explanation. Instead, data suggests that aberrant splicing in the upstream and downstream regions of circRNA producing exons are critical for shaping the circRNAome, and specifically, we observe a conserved impact of missplicing in the immediate upstream region to drive circRNA biogenesis. Collectively, our data show that SFPQ plays an important role in maintaining intron integrity by ensuring accurate splicing of long introns, and disclose novel features governing *Alu*-independent circRNA production.

## Introduction

Gene expression is the output of multiple tightly coupled and controlled steps within the cell, which are highly regulated by a variety of factors and processes. Among these are the physical and functional interactions between the transcriptional and splicing machineries that are of high importance for the generation of both canonical and alternative isoforms of RNA transcripts. One includes a novel class of unique, closed circular RNA (circRNA) molecules.

CircRNAs are evolutionary conserved and display differential expression across cell types, tissues and developmental stages. The highly stable circular conformation is obtained by covalently joining a downstream splice donor to an upstream splice acceptor, a backsplicing process catalyzed by the spliceosome (Memczak *et al*, 2013; Jeck *et al*, 2013; Salzman *et al*, 2013; Ashwal-Fluss *et al*, 2014; Hansen *et al*, 2013). The vast majority of circRNAs derive from coding sequences making their biogenesis compete with the production of linear isoforms (Ashwal-Fluss *et al*, 2014; Salzman *et al*, 2013, 2012). Complementary sequences in the flanking introns can facilitate the production of circRNAs (Dubin *et al*, 1995; Ivanov *et al*, 2015; Jeck *et al*, 2013; Westholm *et al*, 2014; Zhang *et al*, 2014), where the primate-specific *Alu* repeats are found to be significantly enriched in the flanking introns of circRNAs (Jeck *et al*, 2013; Ivanov *et al*, 2015; Venø *et al*, 2015). However, in both human and *Drosophila*, biogenesis of the most abundant and conserved pool of circRNAs tend to be driven by long flanking introns rather than the presence of proximal inverted repeats in the flanking sequences (Westholm *et al*, 2014; Stagsted *et al*, 2019). The biogenesis of circRNAs without inverted repeats is currently not understood in detail, although RNA-binding proteins (RBPs) associating with the flanking introns of circRNAs have been shown to be important (Ashwal-Fluss *et al*, 2014; Conn *et al*, 2015; Errichelli *et al*, 2017).

Here, we aim to identify additional protein factors involved in circRNA biogenesis. To this end, we exploited the enormous eCLIP and RNA sequencing resource available from the ENCODE consortium (ENCODE Project Consortium, 2012). Stratifying eCLIP hits across the genome with circRNA loci coordinates revealed the splicing factor proline/glutamine rich (SFPQ) and Non-POU domain-containing octamer-binding protein (NONO) as highly enriched around circRNAs compared to other exons. Both proteins belong to the multifunctional *Drosophila* behavior/human splicing (DBHS) family with highly conserved RNA recognition motifs (RRMs) (Dong *et al*, 1993) and they are often found as a heterodimeric complex (Knott *et al*, 2016, 2015; Lee *et al*, 2015; Passon *et al*, 2012). The proteins are predominantly located to the nucleus, in particular to the membrane-less condensates known as paraspeckles (Clemson *et al*, 2009; Fox *et al*, 2018), where they play a pivotal role in cellular mechanisms ranging from regulation of transcription by interaction with the C-Terminal Domain (CTD) of RNA polymerase II (Buxadé *et al*, 2008; Rosonina *et al*, 2005; Urban *et al*, 2000), pre-mRNA splicing (Emili *et al*, 2002; Ito *et al*, 2008; Kameoka *et al*, 2004; Peng *et al*, 2002) and 3’end processing (Kaneko *et al*, 2007; Rosonina *et al*, 2005) to nuclear retention (Zhang & Carmichael, 2001) and nuclear export of RNA (Furukawa *et al*, 2015). Recently, SFPQ has been implicated in ensuring proper transcription elongation of neuronal genes (Takeuchi *et al*, 2018) representing an interesting link to circRNAs, as these are highly abundant in neuronal tissues and often derive from neuronal genes (Rybak-Wolf *et al*, 2015).

Here, we show that SFPQ depletion leads to specific deregulation of circRNAs with long flanking introns devoid of proximal inverted *Alu* elements. Moreover, we show that long introns in particular are prone to intron retention and alternative splicing with concomitant premature termination. While premature termination is not the main driver of circRNA deregulation, we provide evidence for a complex interplay between upstream (acting positively on circRNA production) and downstream features (acting negatively) that collectively govern the production of individual circRNAs in the absence of SFPQ. This not only elucidates a conserved role for SFPQ in circRNA regulation but also identifies upstream alternative splicing as an approach towards circRNA production.

## Results

### The DALI circRNAs are defined by long flanking introns and distal inverted Alu elements

To stratify circRNAs by their inverted *Alu*-element dependencies, we characterized the circRNAome in two of the main ENCODE cell lines, HepG2 and K562 (Supplementary Table 1). Using the joint prediction of two circRNA detection algorithms, ciri2 and find_circ, we identified 3044 and 7656 circRNAs in HepG2 and K562, respectively. While proximal inverted *Alu* elements (IAEs) are important for the biogenesis of a subset of circRNAs (Jeck *et al*, 2013; Ivanov *et al*, 2015), we and others have shown that long flanking introns associate with circRNAs loci, particularly for the conserved and abundant circRNAs (Stagsted *et al*, 2019; Westholm *et al*, 2014), and the biogenesis of this group of circRNA species is largely unresolved. To focus our analysis on the non-*Alu*, long intron fraction of circRNAs, we subgrouped circRNAs based on their IAE distance and flanking intron length using median distance and length as cutoffs (Fig. 1A-C), and we observed that these two features show interdependent distributions, where approximately 70% of the top1000 expressed circRNAs group as either Distal-Alu-Long-Intron (DALI) circRNAs or Proximal-Alu-Short-Intron (PASI) circRNAs (Fig. 1D). Apart from long flanking introns and distal IAEs, DALI circRNAs show higher overall expression compared to PASI circRNAs, longer genomic lengths, but a similar distribution of mature lengths (Supplementary Fig. 1A-C). Moreover, almost half of a previously characterized subgroup of circRNAs, the AUG circRNAs (Stagsted *et al*, 2019), derive from DALI circles (Supplementary Fig. 1B), and interestingly, when filtering circRNAs for conservation (in mouse and human), 69-72% of conserved circRNAs are DALI circRNAs (Fig. 1E). This finding suggests that the IAE-dependent biogenesis pathways is not relevant for the most conserved and abundant circRNAs and that other factors must be involved.

**Figure 1:**
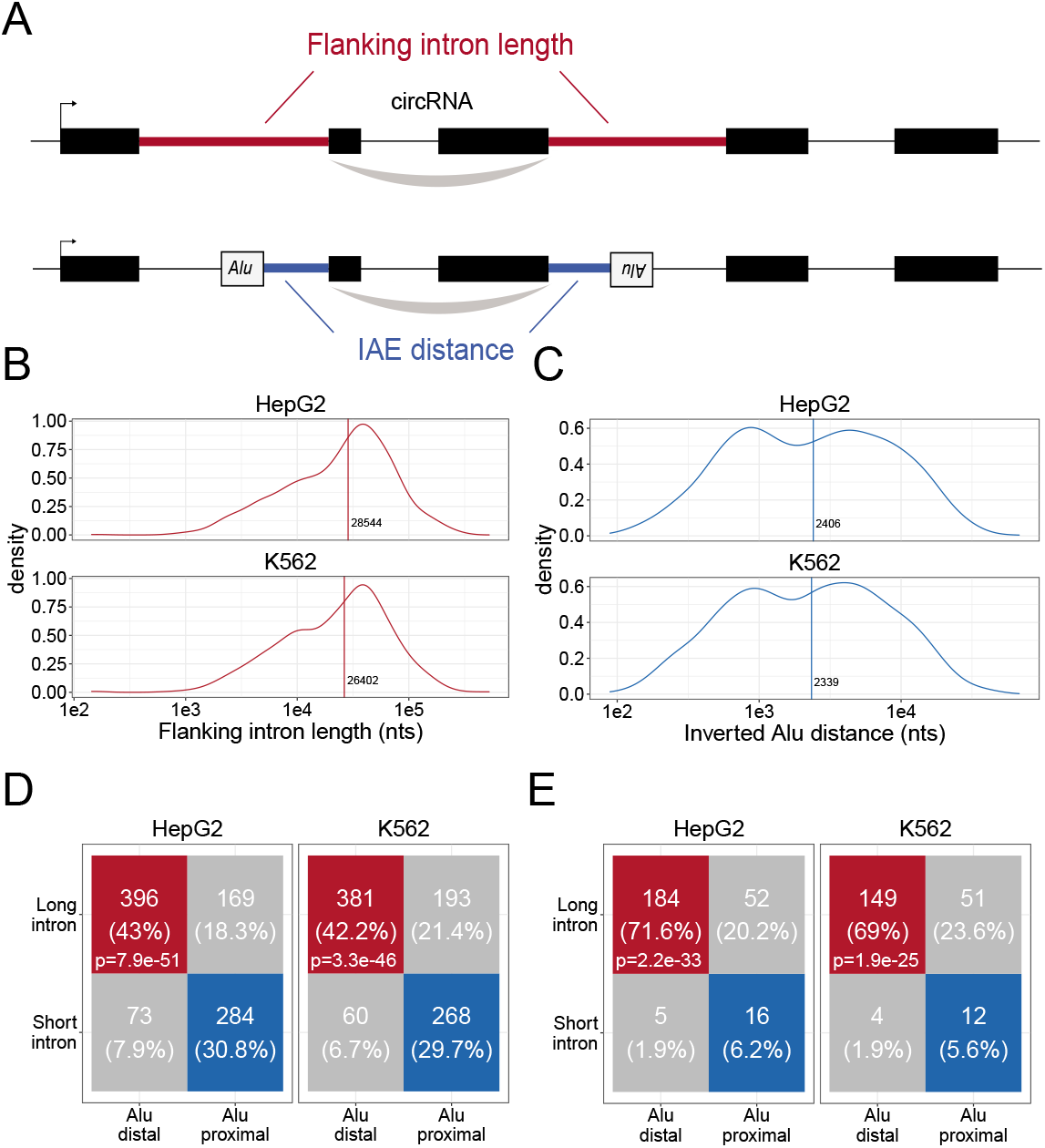
Characteristics of DALI-circRNA. **A)** Schematics showing the flanking intron length (red) defined by the sum of annotated flanking introns and inverted *Alu* element (IAE) distance (blue) defined by the sum of distance to the most proximal IAE. **B-C)** Density plot for the distribution of flanking intron lengths (B) and IAE Distance (C) for the top1000 expressed circRNAs in HepG2 (upper facet) and K562 (lower facet). The vertical line represents the median. **D)** Contingency table showing the 4-way distribution of circRNAs with long and short flanking introns (in respect to the median) and proximal and distal IAEs (also in respect to the median, see B and C) for HepG2 (left facet) and K562 (right facet). The contingency table is color-coded by circRNA subgroup; DALI (distal *Alu*, long flanking introns, in red), PASI (proximal *Alu*, short flanking introns, in blue) and ‘Other’ (unclassified, in grey) circRNAs. The p-values are Fisher’s exact test of independence. **E)** As in D, but for the subset of circRNAs with conserved expression in mouse

### SFPQ and NONO are specifically enriched in the introns flanking DALI circRNAs

In order to identify RNA-binding proteins that could drive circRNA formation, we used the elaborate ENCODE eCLIP data (ENCODE Project Consortium, 2012) (Supplementary Table 2). We scrutinized the immediate flanking regions of the 1000 most highly expressed circRNAs in HepG2 and K562 with the assumption that factors directly involved in backsplicing are likely to bind in the vicinity of the back-splicing sites. We extracted an eCLIP enrichment score using Wilcoxon rank-sum tests between the number of eCLIP reads aligned to circRNA flanking regions (upstream and downstream) compared to flanking regions of host exons, i.e. other exons from the circRNAs expressing genes. We found SFPQ to be the protein most highly enriched for circRNA binding in HepG2 cells, while NONO – a known interaction partner for SFPQ (Dong *et al*, 1993) – was enriched for circRNA binding in K562 cells (Fig. 2A-B, to our knowledge eCLIP datasets on SPFQ in K562 and NONO in HepG2 are not available). Comparing DALI and PASI circRNAs shows that SFPQ is DALI circRNA specific, both upstream and downstream of the circularizing exons (Fig. 2C, p≤1.2e-16), whereas NONO associates with circRNA loci more generally and with the upstream regions of DALI circRNAs specifically (Fig. 2D). SFPQ, like circRNAs, is known to associate with long introns (Iida *et al*, 2020; Takeuchi *et al*, 2018). To exclude that the enrichment seen is a mere bias from the flanking intron length, we extracted a subset of annotated splice acceptor (SA) and splice donor (SD) pairs sampled to match the expression level (linear spliced reads) and flanking intron lengths of DALI circRNAs (denoted ‘DALI-like exons’) (Supplementary Fig. 2A-H). This analysis shows that both SFPQ and NONO were significantly more enriched around circRNA exons compared to sampled DALI-like exons (Supplementary Fig. 2E-H).

**Figure 2:**
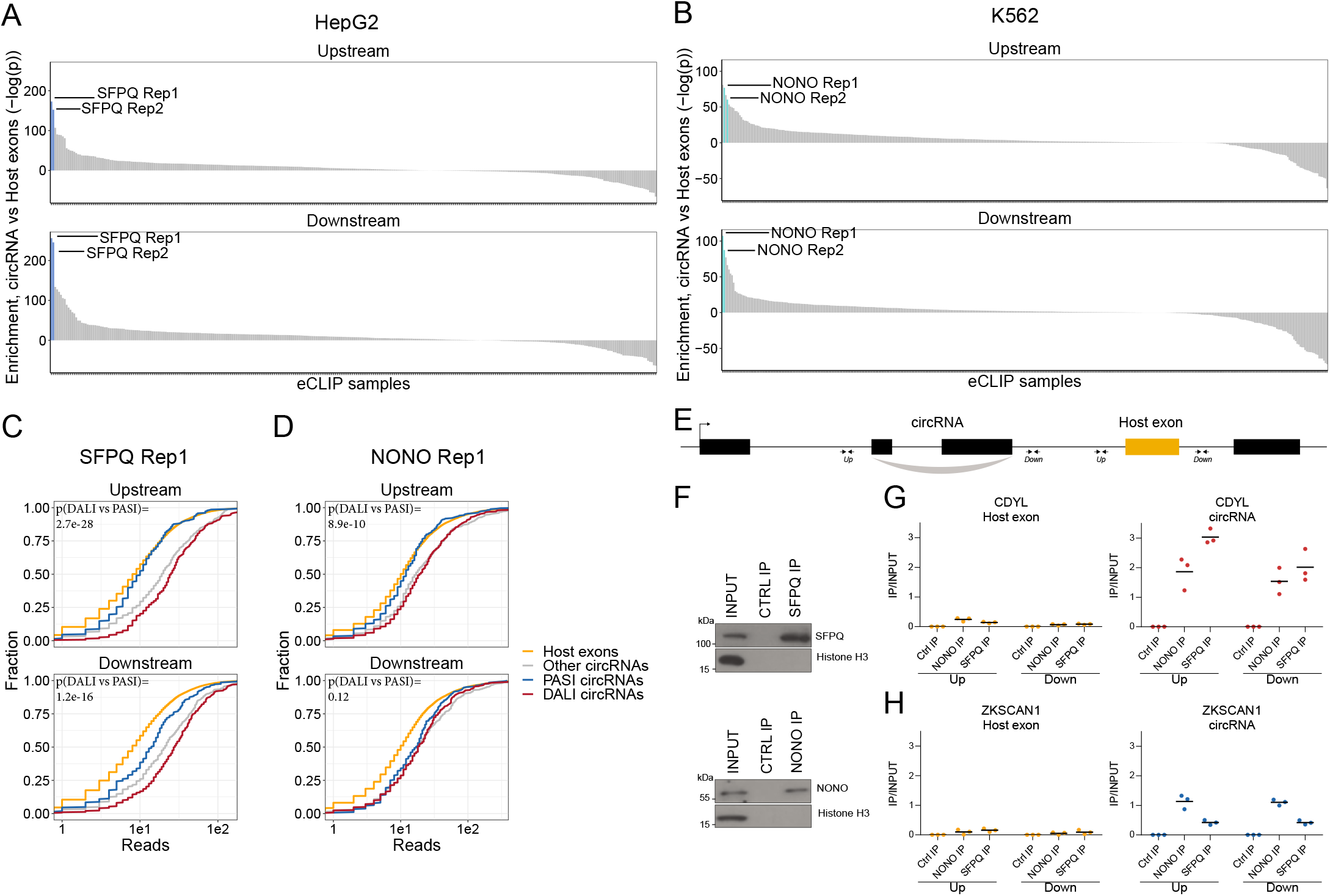
SFPQ and NONO show enriched binding in the flanking regions of DALI circRNAs. **A-B)** Barplot showing enrichment/depletion of eCLIP signal (see Supplementary Table 2) in the vicinity of circRNAs (+/− 2000 nt) compared to host exons (+/− 2000 nt) as determined by Wilcoxon rank-sum tests for HepG2 (A) and K562 (B) eCLIP samples. **C-D**) Cumulative plots of SFPQ (C) and NONO (D) eCLIP read distribution upstream and downstream of circRNA subgroups and host exons as denoted. **E**) Schematic showing localization of primers (+/− 2000 nt) for targeting either upstream (up) or downstream (down) intronic regions of splice sites in respect to circRNA exons or host exon. **F**) Western blotting of immunoprecipitated (IP), endogenous SFPQ or NONO from nuclear fractions of HepG2 cells with Histone H3 as a loading control. **G-H**) qRT-PCR of intronic regions flanking a downstream host gene exon (left facet) or flanking the circRNA producing exon(s) (right facet) of CDYL (G) and ZKSCAN1 (H) upon RNA IP of endogenous SFPQ or NONO from nuclear fractions of HepG2 cells. The relative expression of immunoprecipitate(IP)/input is plotted. Data for three biological replicates are shown.

To validate the binding of SFPQ and NONO on nascent circRNA transcripts, we conducted RNA immunoprecipitation (RIP) qPCR in HepG2 (Fig. 2E-H and Supplementary Fig. 3A-D) and HEK293T cells (Supplementary Fig. 3C-H) and quantified the expression of a panel of representative DALI and PASI circRNAs. Here, the flanking regions of DALI circRNAs, circCDYL and circARHGAP5 (circEYA1 in HEK293T), were significantly enriched for SFPQ binding compared to downstream intronic regions (Fig. 2G-H and Supplementary Fig. 3A, E and G). However, we found no enrichment for PASI circRNAs, circZKSCAN1 (Fig. 3I and Supplementary Fig. 3F) and circNEIL3 (Supplementary Fig. 3B and H). Thus, we conclude that SFPQ and NONO associate with the flanking introns of DALI circRNAs, indicative of a functional role in circRNA biogenesis.

**Figure 3:**
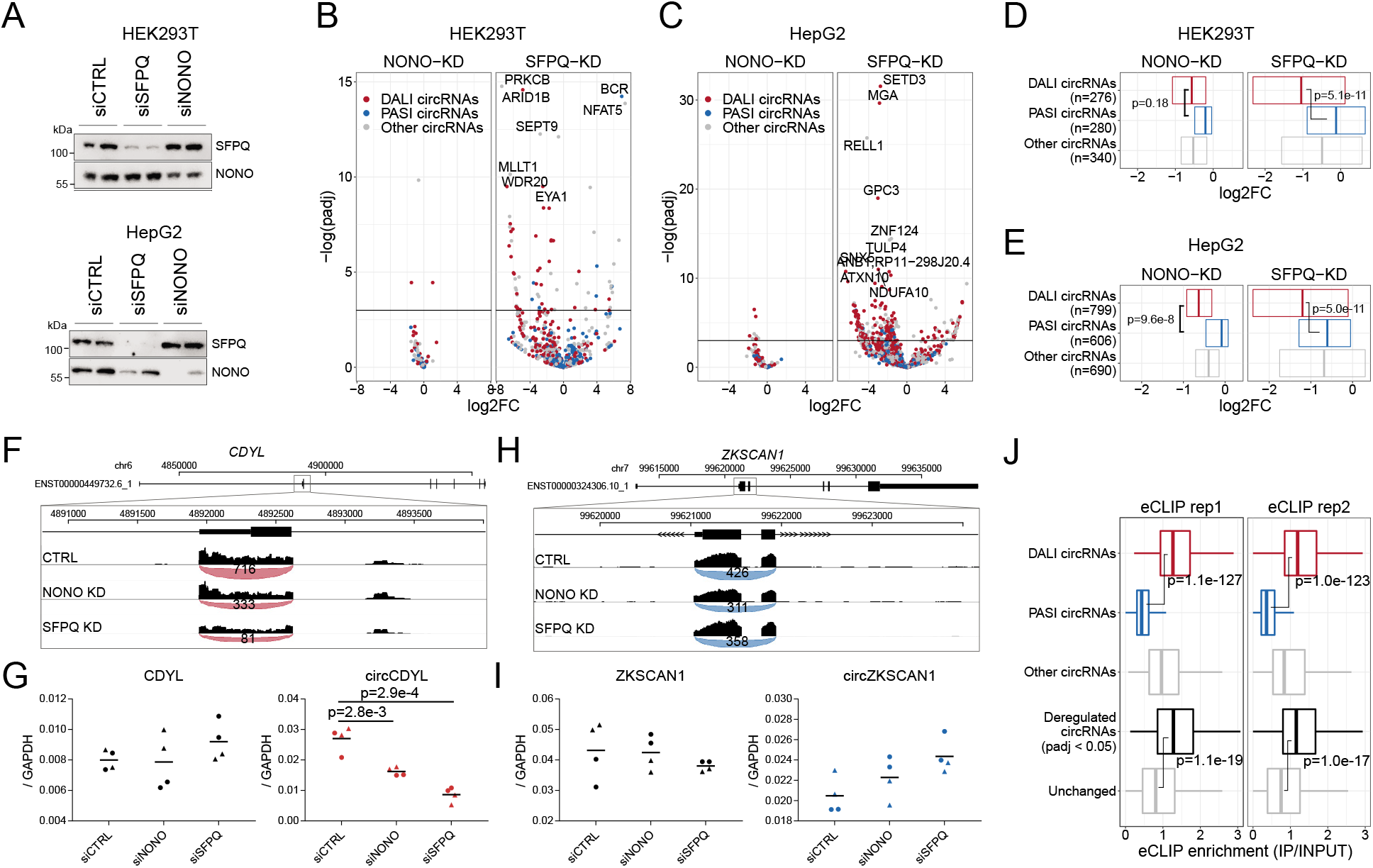
Knockdown of SFPQ affects DALI circRNAs. **A)** Western blotting of proteins from HEK293T (upper panel) and HepG2 (lower panel) cells transfected with either CTRL siRNAs, siRNAs targeting NONO mRNA, or siRNAs targeting SFPQ mRNA using antibodies against SFPQ and NONO as denoted. **B-C)** Volcano plot showing deregulated circRNAs upon NONO (left facet) and SFPQ (right facet) depletion in HEK293T cells (B) or HepG2 cells (C) color-coded by circRNA subgroup; DALI circRNAs (red), PASI circRNAs (blue) and ‘other’ circRNAs (grey). **D-E**) Boxplot showing overall changes in expression (log2Foldchange) of the three circRNA subgroups upon NONO and SFPQ depletion in HEK293T (D) and HepG2 (E) cells. P-values are calculated using two-sided Wilcoxon rank-sum tests. **F)** Genome screen dump of the circCDYL expressing locus with BSJ-spanning reads visualized as junction-track in the IGV browser **(G)** qRT-PCR quantification of circCDYL and linear CDYL expression upon SFPQ and NONO-depletion in HepG2 cells relative to GAPDH mRNA using two different siRNA designs for each target. Data for four biological replicates are shown. P-values are calculated using student’s two-tailed t-test. **H-I)** as in F and G, but for the PASI circRNA, circZKSCAN1. **J)** Boxplot showing eCLIP reads for SFPQ flanking the three circRNA subgroups as well as the significantly deregulated circRNAs (with padj < 0.05) compared to the unchanged in as observed in SFPQ-depleted HepG2 cells. P-values are calculated using two-sided Wilcoxon rank-sum tests.

### SFPQ and NONO depletion represses DALI circRNAs production

To study the impact of SFPQ and NONO on circRNA production, we depleted SFPQ and NONO in HepG2 and HEK293T cells using two different siRNAs for each target (Supplementary Fig. 4A, Supplementary Tables 3 and 4). Western blot and RT-qPCR (Fig. 3A, Supplementary Fig. 4B-E) showed that expression of both proteins was efficiently reduced upon siRNA treatment, although, unexpectedly, the expression levels of *SFPQ* mRNA appeared unaffected by SFPQ knockdown and greatly elevated upon NONO depletion (Supplementary Fig. 4C and E). This, we speculate, is the result of compensatory effects or autoregulatory mechanisms. We performed total RNA sequencing of the knockdown samples, and conducted gene expression analysis of circRNA and mRNAs. Principal Component Analysis (PCA) of HEK293T and HepG2 samples shows clear grouping of treatments (SFPQ, NONO and CTRL), both on mRNA and circRNA levels (Supplementary Fig. 4F-I), suggesting that most of the variance between samples are explained by the knockdown. Although for HepG2 two samples (siSFPQ1_rep1 and siNONO2_rep2) display outlier signatures and were thus removed in downstream analyses.

The differential circRNA expression analysis showed that DALI circRNAs are generally reduced upon SFPQ depletion, whereas PASI circRNAs are practically unaffected in both HEK293T (Fig. 3B and D) and HepG2 (Fig. 3C and E) cells. For NONO, we observed almost no impact on circRNAs production in both cell lines (Fig. 3B-E). This could either indicate that NONO is less involved in circRNAs biogenesis, or that the effect is in part masked by the concomitant upregulation of SFPQ observed upon NONO depletion. Consistently, RT-qPCR analyses of abundant DALI circRNAs, circCDYL (Fig. 3F, Supplementary Fig. 5C), circARHGAP5 (Supplementary Fig. 5A) and circEYA1 (Supplementary Fig. 5E), and PASI circRNAs, circZKSCAN1 (Fig. 3G, Supplementary Fig. 5D) and circNEIL3 (Supplementary Fig. 5B and F) confirmed repressed expression of DALI circRNAs and unchanged PASI circRNAs expression relative to host gene levels. Finally, to support a direct role for SFPQ in circRNA formation, we overlaid the results from SFPQ-depleted HepG2 cells with the SFPQ eCLIP data and observed a significant association between SFPQ binding in the flanking regions of DALI circRNAs, as expected, but also a clear association with deregulated circRNAs compared to unchanged circRNAs (Fig. 3J, p < 1e-17, Wilcoxon rank-sum tests). In addition, we examined previously published total RNAseq from SFPQ conditional knock-out (KO) mouse brain (Takeuchi *et al*, 2018) (Supplementary Table 5). Here, as in human cell lines, DALI and PASI circRNA are prevalent subclasses (Supplementary Fig. 6A) with DALI circRNAs showing higher abundancy compared to PASI circRNAs (Supplementary Fig. 6B). SFPQ depletion in mouse brain affects global circRNA expression (Supplementary Fig. 6C); however, in contrast to HEK293T and HepG2 cells, we found a more equal distribution of up- and downregulated circRNAs upon SFPQ removal (Supplementary Fig. 6D-E), and we detect a clear tendency for DALI circRNAs to be more prone to SFPQ-mediated regulation (25% vs 5% showing significant deregulation, Supplementary Fig. 6F, p=8.2e-80, Fisher’s exact test). Collectively, these findings suggest that SFPQ (and to a lesser degree NONO) regulates DALI circRNA biogenesis.

### SFPQ depletion affects alternative splicing and intron retention in long genes

Next, to understand the impact of SFPQ and NONO on transcription and splicing in general, we used the RNAseq data to investigate SFPQ/NONO-sensitive mRNAs. Here, we found that SFPQ-depletion triggers a general repression of long genes (stratified by median gene length, Fig. 4A). The read distribution of highly repressed genes showed a peculiar expression profile with unaffected read densities at the genic 5’ends but with dramatic reduction at the 3’end in HepG2 cells (Supplementary Fig. 7A-D) indicating that the transcription machinery drops off mid gene. This prompted us to survey genes globally for a ‘drop-off’ phenotype. Thus, we subgrouped genes into their expression profile by slicing each gene into 20 equally sized bins and conducting differential gene expression on all bins. To subgroup genes with of similar profiles in an unsupervised manner, we clustered the log2foldchanges across genes into five categories, denoted kc1-5, using k-means clustering (Fig. 4B). Here, kc5 but also kc3 and 4 showed ‘drop-off’ effects but to different degrees, and interestingly, the effect correlates with gene length (Fig. 4B-C). We obtain almost identical results from SFPQ-depleted HEK293T cells (Supplementary Fig. 8A-C) and mouse brain (Supplementary Fig. 8F-H).

**Figure 4:**
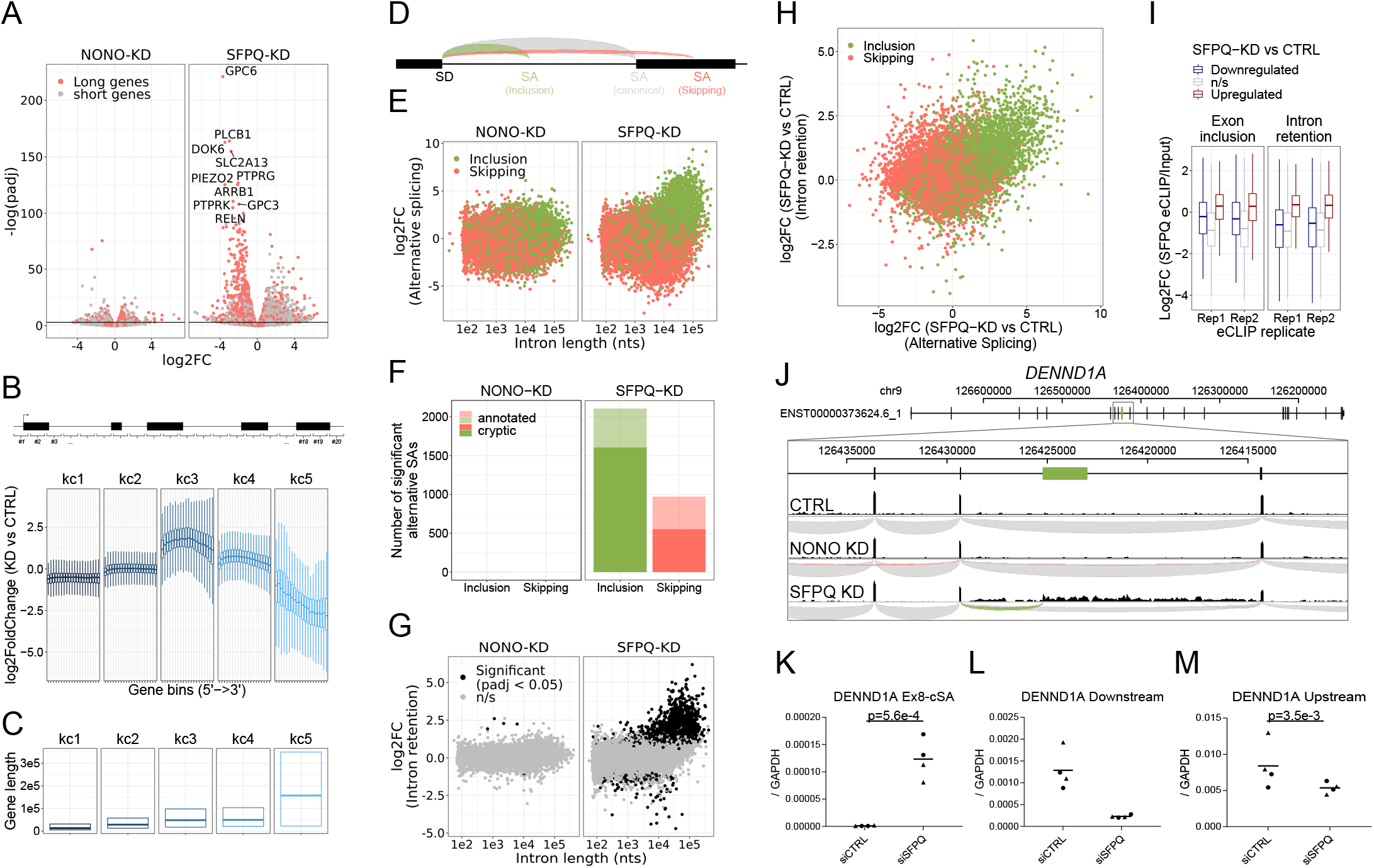
SFPQ ensures long-gene expression and suppresses cryptic splicing. **A)** Volcano plot depicting differential expression of annotated genes upon NONO or SFPQ KD compared to CTRL in HepG2 cells, stratified by median gene length into ‘long’ and ‘short’ genes as denoted. **B)** Boxplot showing binned expression of clustered genes. Each gene is sliced into 20 equally sized bins, and the differential expression of each bin is determined and subgrouped into five k-means clusters (kc) (see Methods). **C)** Boxplot showing gene lengths distribution (.25,.5 and.75 quantiles) stratified by clusters obtained in B. **D)** Schematic representation of alternative splicing, where canonical (grey) denoted the most abundant splicing from the splice donor in question. Inclusion (green) and skipping (red) denotes an alternative splicing event shorter or longer than canonical, respectively**. E)** Scatter plot showing alternative splicing in NONO and SFPQ depleted samples as a function of canonical intron length and color-coded by type of splicing (either inclusion or skipping, see schematics in D. **F)** Barplot with the number of unique alternative splicing events showing significant deregulation upon NONO and SFPQ depletion stratified by inclusion(green) and skipping (red), and whether the alternative SA site is annotated (transparent) or not (opaque). **G)** Scatter plot showing effects on intron retention (IR) upon SFPQ and NONO depletion as a function of intron length, color-coded by significance (adjusted p-value < 0.05) as denoted. **H)** Scatterplot showing for each detectable intron the correlation between changes in exon-inclusion/skipping (red/green) and intron retention upon SFPQ depletion. **I)** Boxplot showing the IP/Input enrichment of SFPQ eCLIP reads in introns harbouring an exon inclusion or an intron retention event color-coded by whether the event is up or down (red or blue, respectively) or not significant (n/s, grey). **J)** Schematic showing coordinates and full genic locus of *DENND1A* (top panel) and exon 8 and 9 with alternative, unannotated exon in-between (green, middle panel). Merged intron-spanning reads (lower panel) from CTRL, NONO-KD and SFPQ-KD samples (HepG2) are shown and color-coded by splicing type; canonical (grey), inclusion (green) and skipping (red), see D. **K-M)** qRT-PCR analysis of alternative splicing event (K), upstream expression (L) and downstream expression (M) relative to *GAPDH* mRNA. Data for four biological replicates are shown. P-values are calculated using student’s two-tailed t-test.

Upon inspection of the downregulated genes in our samples, we found an upregulation of alternative splicing in the SFPQ KD samples (Supplementary Fig. 7). We classified all alternative splicing events as either inclusion or skipping relative to their respective canonical isoform (Fig. 4D) and performed differential expression analysis using DESeq2. This showed an extensive change (mostly upregulation) of alternative splicing events correlating with intron length in both HepG2, HEK293T and mouse brain (Fig. 4E, Supplementary Fig. 8D and I). Of the 2106 significantly deregulated inclusion events in HepG2, more than 96% are upregulated and of these, 76% are not annotated by gencode (Fig. 4F, in HEK293T: 95% upregulated, 78% unannotated, in mouse: 90% upregulated, 88% unannotated: data not shown), and consequently, we suggest that these events are mostly cryptic or aberrant splicing. Furthermore, analyzing the levels of intron retention by quantifying unspliced intronic reads shows a very similar intron-length-dependent pattern with significant retention of long intron upon SFPQ depletion (Fig. 4G, Supplementary Fig. 8E and J). Consistently, we find a clear correlation between exon inclusion and intron retention (Fig. 4H), and a clear enrichment of SFPQ eCLIP signal in regions subjected to alternative splicing and intron retention (Fig. 4I). As an example, for *DENND1A*, we observe a previously unannotated splicing event joining exon eight to an alternative splice acceptor dinucleotide (AG) residing in intron eight of this gene (Fig. 4J), which is only detectable upon SFPQ knockdown (Fig. 4K), and, in *DENND1A*, this cryptic event marks the transition from unaffected to repressed state, as quantification of the upstream region shows modest to no effect between control and knockdown, whereas the downstream region is highly suppressed (Fig. 4K-M).

Collectively, this suggests that intron retention and alternative splicing are conserved effects of SFPQ depletion, and that SFPQ plays a vital role in splicing integrity for long introns in particular.

### SFPQ depletion results in premature termination events

In order for alternative splicing to result in premature termination of transcription, the alternative/cryptic-included exons need to harbor a polyA-signal that can serve as a functional terminator of transcription. To investigate the magnitude of polyA-signal appearance in SFPQ knockdown samples, we subjected SFPQ and NONO-depleted HEK293T cells to 3’end quantSeq (Supplementary Fig. 9A, Supplementary Table 6).

Putative polyA-signals were retrieved using the MACS2 callpeak algorithm, and to further increase the signal to noise ratio, we characterized each peak by the presence of a bonafide polyA-signal (PAS: AAUAAA or AUUAAA). Furthermore, for each quantseq peak, we also extracted the highest prevalence of A’s in all possible 15-nucleotide windows to reduce non-polyA-tail artefacts in the samples. The fraction of PAS-containing peaks dropped markedly when regions with 14 or 15 nucleotides A-stretches were found (Supplementary Fig. 9B), suggesting that these A-rich peaks are likely polyA-tail-independent artefacts and were thus removed from the analysis. The remaining peaks were classified as PAS sites, and for all PASs, the genic origin was annotated, and the differential usage was determined by DESeq2. This showed a clear enrichment on intronic PAS and a repression of exonic PAS usage upon SFPQ knockdown (Fig. 5A). As before, NONO-depletion only showed a modest effect.

**Figure 5:**
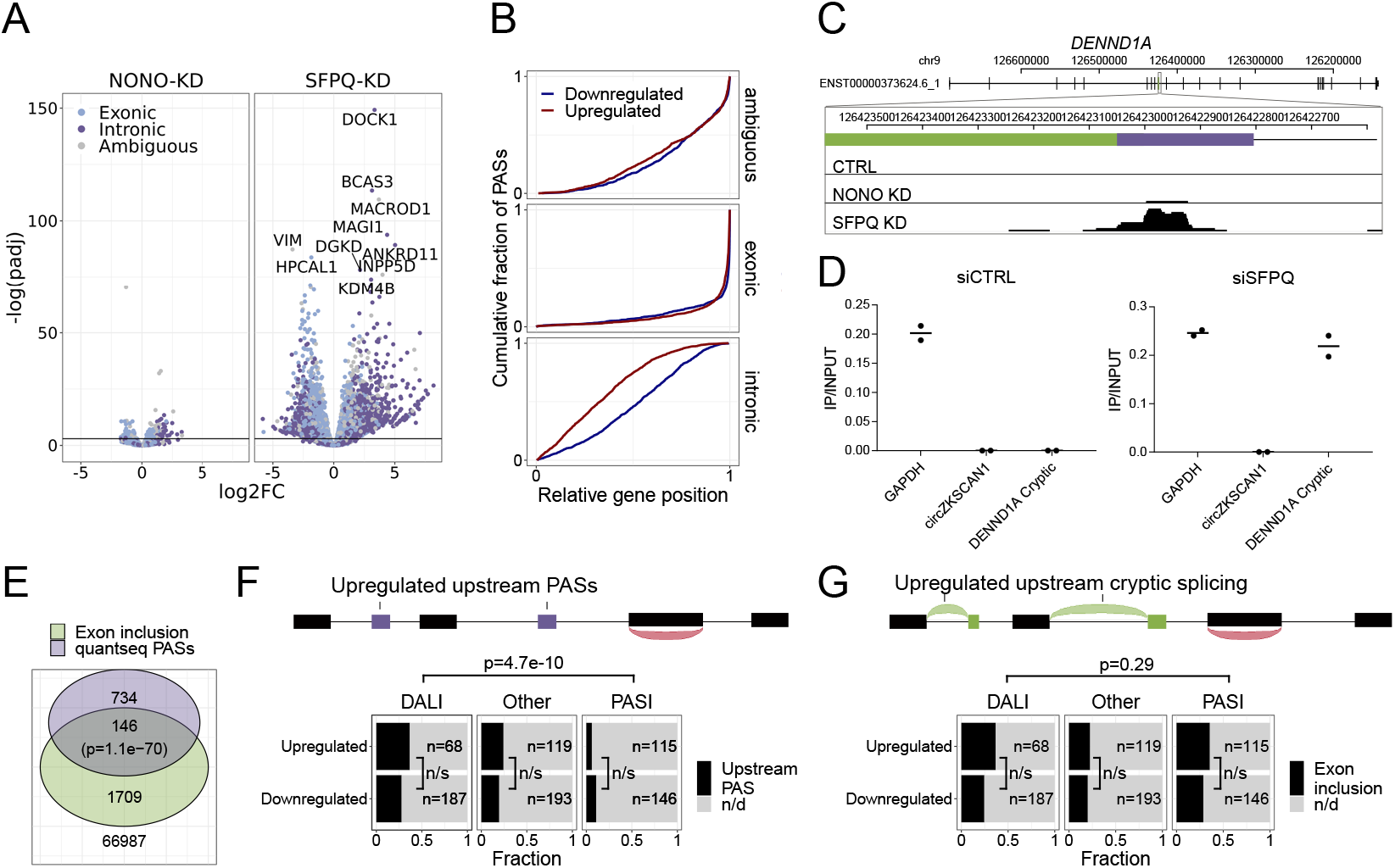
SFPQ depletion activates intronic polyA signal and premature termination. **A)** Volcano plot showing deregulated PAS usage as measured by quantseq upon NONO and SFPQ depletion in HEK293T cells. PAS signals are color-coded by their genic origin; intronic (dark blue), exonic (light blue) or ambiguous (grey). **B)** Plot showing the cumulative fraction of PASs as a function of relative genic position stratified by genic origin (ambiguous, exonic or intronic, vertical facets) and color-code by whether the PAS is significantly up (red) or downregulated (blue) upon SFPQ knockdown. **C**) Schematic representation of the DENND1A exon 8-9 locus with alternative exon (green) and putative PAS element (purple). Below, merged quantseq coverage from each experiment. **D**) qRT-PCR on input and oligo-dT purified RNA from control and SFPQ-depleted HEK293T cells using amplicons specific for GAPDH (positive control), circZKSCAN1 (negative control) and the alternative SFPQ-activated exon. Values reflect ratios between oligo-dT purified and input quantities. Data for two biological replicates are shown. **E**) Venn diagrams showing the number of unique introns with co-occurring upregulation of PAS and upregulated alternative splicing. The number of expressed introns without any evidence of enriched PASs or alternative splicing is denoted below the diagram. P-values are calculated by Fisher’s exact test. **F-G**) Schematic showing the outline of the analysis (upper panel): For each circRNA, the locus spanning from the promoter to the circRNA splice donor was interrogated for the presence of quantseq PASs (F) or exon inclusion (G). Barplot (lower panel) showing the fraction of upregulated and downregulated circRNAs upon SFPQ depletion in HEK293T cells with evidence of a concomitant upregulated upstream PAS (F) or an upstream exon inclusion event (G). Numbers indicate the total number of circRNAs in each group. P-values are calculated by Fisher’s exact test.

As an upstream termination impacts downstream elements, we determined the relative genic position of up- and downregulated PASs. This showed a clear and general 5’region tendency of upregulated vs downregulated intronic PASs (Fig. 5B), suggesting that activation of upstream PASs may subsequently repress the usage of downstream PASs. In addition, activation of upstream PASs were particularly pronounced in kc4 and 5 (Supplementary Fig. 9C) indicating that the ‘drop-off’-phenotype may be a consequence of intronic PAS activation and premature termination.

To investigate how alternative splicing relates to premature termination in a global manner, we assessed for the co-presence of alternative exon inclusion and significantly enriched PASs across all expressed introns (Fig. 5E). Overall, this showed a significant overlap (Fig. 5E) with kc4 exhibiting the highest degree of overlap with 72 distinct introns harboring both events (Supplementary Fig. 9D). For DENND1A, where cryptic splicing marks the transition from unaffected to repressed state, we also observe a clear PAS with a consensus polyA signal (Fig. 5C). This was validated using polyA enrichment, where the alternative transcript is oligo-dT purified as effectively as *GAPDH* only upon SFPQ knockdown. Collectively, this suggests that a notable fraction of genes exhibit alternative splicing and premature termination upon SFPQ knockdown with increased probability for longer introns, underscoring, once again, the importance of SFPQ in gene expression.

### circRNA deregulation is not explained by premature termination

If SFPQ depletion results in wide-spread increase in premature termination, the observed deregulation of circRNAs in our dataset could simply be explained by incomplete transcription and not as a biogenesis effect *per se*. This notion is consistent with the fact that circRNAs in general and DALI circRNAs in particular associate with long flanking introns prone to alternative splicing and premature termination. However, not all circRNAs were depleted upon SFPQ knockdown and particularly in mouse brain, the DALI circRNAs were affected in both directions (i.e. up- and downregulated). To test whether the deregulation of circRNAs is driven by premature termination, we stratified circRNAs by their host gene clusters. This showed that while most circRNAs derive from kc1, 2 and 4, roughly the same expression profile is observed across all clusters (Supplementary Fig. 10A-F). In addition, comparing backsplicing to linear splicing from the circRNA producing loci, no clear correlation was observed, suggesting that the circRNA deregulation is not a mere consequence of transcription levels (Supplementary Fig. 10G-I). Finally, counting the prevalence of upstream (from the SD) significant intronic quantseq PASs (Fig. 5F) or alternative splicing events (Fig. 5G) there is no significant difference between up- and downregulated circRNAs, and for alternative splicing no difference between DALI and PASI circRNAs, whereas premature termination is more prominent upstream of DALI circRNAs. Collectively, we argue that premature termination is not the main driver of circRNA deregulation.

### Extracting features important for circRNA biogenesis

But what is then the underlying explanation for the deregulated expression of DALI circRNAs upon SFPQ depletion? As no single feature captures the circRNA deregulation accurately, we turned to multivariate regression analysis. Here, we collected a number of genic features (up- and downstream intron lengths, IAE distance, annotated distance to promoter and termination, and genomic length of circRNA), and differential expression data upon SFPQ depletion (linear up- and downstream splicing, flanking alternative splicing, upstream alternative splicing, up- and downstream intron retention) (Fig. 6A). Pairwise correlation of all features shows modest redundancy but for certain combinations, such as 5’ linear splice (5’S) and 3’ linear splicing (3’S), we find a high level of positive interdependence (Supplementary Fig. 11 and 12), whereas intron retention generally correlates negatively with linear splicing (5’IR vs 5’S and 3’IR vs 3’S). In fact, linear splicing correlates negatively with all other features included in both HepG2 cells and mouse brain (Supplementary Fig. 11 and 12).

**Figure 6:**
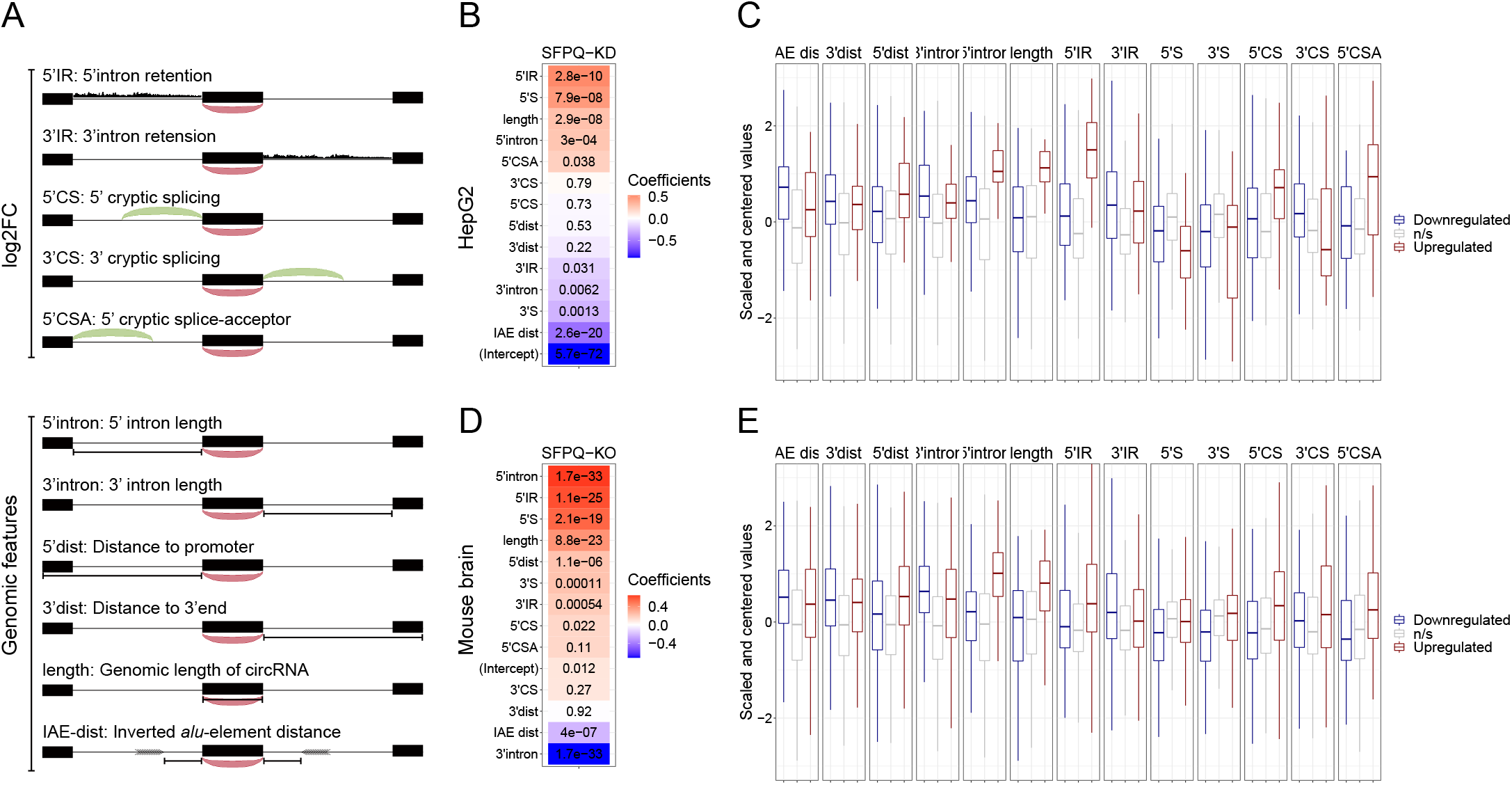
Multiple features contribute to circRNA regulation by SFPQ. **A**) Schematic representation of features used in analysis. **B**) Heatmap showing the feature coefficients from modelling circRNA deregulation (log2FoldChange) upon SFPQ depletion in HepG2 cells. The numbers within the heatmap are the associated p-values. **C**) Boxplot showing the centered and scaled feature-values for significant up (red), significant down (blue), and unchanged (grey) circRNAs in HepG2. **D-E)** as in B and C using mouse brain data.

Splitting the quantified circRNAs into train and test sets (80:20 ratio), we trained a generalized linear model (GLM) against the observed circRNA log2foldchange. As all features were standardized, the resulting coefficients serve as a proxy for feature importance. Here, ranking features by coefficient, it is evident for both HepG2 and mouse brain that 5’ features generally correlate positively with circRNA production, whereas 3’ features (and IAE distance) correlate negatively (Fig. 6B and D). As seen in both HepG2 and mouse brain, certain features, such as 5’IR (upstream intron retention), 5’ intron (upstream intron length) and 5’CSA (upstream cryptic SA usage), are highly distinctive for upregulated circRNA (Fig. 6C and E). Interestingly, features showing high interdependence have opposing effects on circRNA abundance. Here, according to the model, in HepG2, 5’S and 3’S impose positive and negative impact on model prediction, respectively (Fig 6B), whereas features showing anti-correlation (5’S and 5’IR) are both ascribed a positive coefficient in both HepG2 and mouse (Fig. 6B and D). This was also observed in HEK293T, although less convincingly partly due to low sequencing depth in these samples (Supplementary Fig 13A-D). Also, while the performance of the model on the test-set is modest but significant (Supplementary Fig. 14A and C, Pearson correlations: 0.38 (HepG2, p=2.3e-13) and 0.39 (mouse brain, p=6.0e-41), we observe convergens between the HepG2 and mouse brain-derived coefficients suggesting that the obtained features are conserved aspects of SFPQ-mediated circRNA regulation. Here, the most notable difference between HepG2 and mouse brain is the estimated intercept term (Supplementary Fig. 14C), which we interpret as the difference in cellular context, suggesting that the overall impact of SFPQ on circRNA expression may depend on other unidentified factors. Conclusively, the aberrant splicing and intron retention caused by SFPQ depletion impacts circRNA production and explain in part the DALI-specific deregulation observed.

## Discussion

The biogenesis of circRNAs is typically ascribed the presence of proximal inverted repeat elements positioning the two splice sites involved in backsplicing into close proximity. For example, *Alu* elements are frequently found close to circRNA exons (Ivanov et al., 2015b; Jeck et al., 2013), but since they are primate-specific (Lander et al., 2001), the biogenesis of most highly expressed and conserved circRNAs can not be explained by the presence of such elements (Stagsted *et al*, 2019). Additionally, by RNA association and dimerization, RNA-binding proteins have a similar ability to juxtapose splice sites destined for backsplicing, although this currently only seems to apply to a few specific cases (Conn *et al*, 2015; Errichelli *et al*, 2017; Ashwal-Fluss *et al*, 2014; Kramer *et al*, 2015). Here, we attempted to further disclose the impact of RBPs on circRNAs biogenesis and to reveal features important for backsplicing. First, we subgrouped circRNAs into the *Alu*-dependent subset characterized by proximal IAEs and short flanking introns, termed PASI circRNAs, and the *Alu*-independent circRNAs with distal IAEs co-occurring with long flanking introns, the DALI circRNAs. By utilizing the extensive eCLIP resource made available by the ENCODE consortium, we identified SFPQ and NONO as two potentially interesting candidates, both associating significantly with the DALI circRNA producing loci. In both HepG2 and HEK293T cell lines as well as in mouse brain conditional knockouts, we observed a general deregulation of DALI circRNAs upon depletion of SFPQ and only subtle effects upon NONO KD, possibly due to the concomitant upregulation of SFPQ in these samples. Thus, we mostly focused our analysis on SFPQ. Here, apart from dramatic changes in DALI circRNA expression, we observed across all samples (HepG2, HEK293T and mouse brain), that the absence of SFPQ results in aberrant splicing with extensive induction of cryptic splice acceptor sites, particularly in long introns. This correlates with a similar increase in intron retention, suggesting that these two phenotypes are closely coupled. Consistently, recent studies have shown SFPQ to associate with long introns (Iida *et al*, 2020; Takeuchi *et al*, 2018) and to be vital in regulating alternative splicing of long target genes ultimately affecting neural differentiation (Luisier *et al*, 2018) and axon development (Thomas-Jinu *et al*, 2017). Interestingly, circRNAs tend to originate from longer genes, especially neuronal genes (Ragan *et al*, 2019; Rybak-Wolf *et al*, 2015; Szabo *et al*, 2015; You *et al*, 2015), and exons prone to circularize are more frequently flanked by longer introns than non-circularized exons (Jeck *et al*, 2013), supporting SPFQ-sensitive regulation of circRNAs, in particular DALI circRNAs.

Using quantseq analysis, we found SFPQ-depletion to activate the use of intronic polyA-sites, which in many cases overlap with aberrant cryptic splicing thereby resulting in premature termination. Consistently, we also observe decreasing signal across the gene body in the absence of SFPQ. A model in which SFPQ facilitates the recruitment of CDK9 to the CTD of RNA polymerase II to maintain transcription elongation was recently proposed to explain the ‘drop-off’ effect seen upon SFPQ depletion (Takeuchi *et al*, 2018; Hosokawa *et al*, 2019). Instead, we claim that this is partly explained by the induced cryptic splicing and subsequent premature termination, emphasizing that transcription is highly coupled to splicing. SFPQ was initially found to be associated with the polypyrimidine tract, aid in the assembly of the spliceosome and be critical for the second catalytic step in splicing (Ajuh *et al*, 2000; Makarov *et al*, 2002; Gozani *et al*, 1994; Patton *et al*, 1993), supporting its role in splicing fidelity.

While DALI circRNAs in HepG2, HEK293T, and mouse brain are generally sensitive to SFPQ depletion, premature transcription termination fails to explain the observed circRNA levels. Instead, upstream intron length and aberrant splicing in the immediate upstream region have stimulating effects on circRNAs biogenesis, while, although less clear, cryptic events downstream show more detrimental impact. We speculate that SFPQ plays an imperative role in splicing fidelity, a role that becomes increasingly important with intron length. In the absence of SFPQ, by less recruitment to the RNA polymerase II by e.g. Dido3 (Mora Gallardo *et al*, 2019) or due to increased sequestering in paraspeckles, splicing malfunctions resulting in intron retention and reduced backsplicing. With the persistent presence of intronic sequences, cryptic and aberrant splicing become more likely, and cryptic exon inclusions in AU-rich introns will in many cases contain PAS-signal and thus cause premature termination. Consistent with this, RNA polymerase has been shown to be stalled at AT-rich sequences (Henriques *et al*, 2013; Palangat *et al*, 2004), thus allowing a window of opportunity for splicing- and possible cleavage-directed transcription termination.

For circRNAs, the splice-sites involved in backsplicing must be protected from linear splicing as backsplicing occurs less effectively than canonical splicing. In particular, the SA has to remain unspliced until the RNA polymerase reaches the downstream SD. This can be facilitated by a fast polymerase elongation rate (Zhang *et al*, 2016) or by the lack of spliceosomal components (Liang *et al*, 2017). While SFPQ-depletion generally induces cryptic splicing imposing additional splice site competition, it also potentially eliminates upstream linear splicing and thus uncouples the SA from any upstream SD. This potentially exposes the upstream SA for backsplicing consistent with the observed positive impact of cryptic events (5’ intron retention and 5’cryptic SA). In addition, we also observe that the mere length of the 5’ intron has an important predictive value for circRNA formation. This, we hypothesize, is due to the high correlation with aberrant splicing and intron retention; both features are limited to detection in RNAseq. Supposedly, many of the cryptic events are not detectable in a steady state sequencing approach as they are likely unstable and subjected to nuclear quality control or nonsense mediated decay (NMD), and therefore intron length may serve as a useful proxy for aberrant splicing upon SFPQ-depletion. Furthermore, SFPQ is often described in various protein complexes with some comprising FUS and the nuclear resolvase DHX9. Like SFPQ, FUS has been shown to act in various processes within the cell, such as transcription regulation and RNA metabolism (Lagier-Tourenne *et al*, 2012; Kwiatkowski *et al*, 2009; Vance *et al*, 2009) but also to associate with the 5’ss of long introns (Nakaya *et al*, 2013; Lagier-Tourenne *et al*, 2012), especially those flanking circularizing exons, and hereby regulate circRNA biogenesis (Errichelli *et al*, 2017). DHX9 has, on the other hand, been shown to unwind intronic base pairing and thereby reduce the production of *Alu*-dependent circRNAs (Aktaş *et al*, 2017; Errichelli *et al*, 2017). Both proteins show interesting circRNA regulation abilities which could act cooperatively with SFPQ and thus affect the fate of DALI circRNAs upon SFPQ depletion.

Conclusively, we show that SFPQ is a key regulator of DALI circRNAs production by controlling and enforcing accurate long intron splicing. This highlights the complex and intricate relationship between splicing in general and backsplicing in particular. Furthermore, SFPQ has been associated to diverse neurological diseases, such as ALS (Thomas-Jinu *et al*, 2017; Luisier *et al*, 2018) and FTLD (Ishigaki *et al*, 2017), and may prove to be a critical for maintaining the circRNAome in these and other neurodegenerative pathologies. And while in steady state scenarios, cryptic splicing is negligible, it is interesting to speculate whether upstream cryptic splicing is generally involved in DALI circRNA production providing a useful tool for manipulating circRNA production without impacting host gene expression.

## Materials and Methods

### Cell lines and transfections

HEK293T cells (Invitrogen), HepG2 (ATCC) were cultured in Dulbecco’s modified Eagle’s media (DMEM) with GlutaMAX (Thermo Fisher Scientific) supplemented with 10% foetal bovine serum (FBS) and 1% penicillin/streptomycin sulphate (P/S). All cells were kept at 37°C in a humidified chamber with 5% CO2. Knockdown of SFPQ and NONO were carried out using transient transfections of siRNAs accordingly (siRNA sequences in Supplementary Table 7): For HEK293T, approximately 250.000 cells were plated in a 6-well dish and transfected with a final concentration of 22.5 nM siRNA using siLentFect Lipid Reagent (Bio-Rad) accordingly to manufacturer’s protocol. 48 hours post transfection, the cells were replenished with new media and re-transfected using Lipofectamine 2000 (Thermo Fisher Scientific). After additional 48 hrs, cells were harvested. For HepG2, approximately 400.000 cells were plated in a 6-well dish and reverse transfected with a final siRNA concentration of 50 nM using Lipofectamine RNAiMax (Thermo Fisher Scientific). 48 hrs post transfection, the cells were trypsinized before reverse transfected for the second hit using Lipofectamine 2000 (Thermo Fisher Scientific). After 48 hrs, the cells were harvested (see Supplementary Fig. 4A for experimental outline). For all experiments, cells were harvested by washing in 1xPBS and subsequent centrifugation at 1200 rpm at 4°C for 4 min. 66.6% of the harvested cells was used for RNA isolation, which was carried out using TRIzol^®^ Reagent (Thermo Fisher Scientific) accordingly to manufacturer’s protocol. Except for RNA used for RNAseq and RIP, one μg RNA was subjected to DNase I treatment (Thermo Fisher Scientific #EN0521) accordingly standard protocol prior to subsequent analysis. The remaining cells (33.3%) were used for protein isolation; after centrifugation, the cell pellets were resuspended in 2xSDS loading buffer [125 mM Tris–HCl pH 6.8, 20% glycerol, 5% SDS, and 0.2 M DTT] and boiled at 95°C for 5 min.

### RT-PCR and RT-qPCR

One μg of DNase-treated total RNA was reverse transcribed using the M-MLV Reverse Transcriptase kit (Thermo Fisher Scientific) according to manufacturer’s protocol with the use of random hexamers to prime the reaction. In case of RT-PCR, the reaction was conducted with 30 cycles of PCR with or without RT enzyme (Primers listed in Supplementary Table 7). The products were visualized by 1% agarose gel electrophoresis and verified using Sanger sequencing. For quantitative PCR, cDNA was mixed with Platinum^®^ SYBR^®^ Green I Master kit (Invitrogen) and ran on Light cycler 480 II instrument (Roche). The reactions were carried out in technical triplicates. The obtained Ct values for each triplicate were transformed (2-Ct) and averaged (σ). All samples were normalized to GAPDH. The results were visualized using GraphPad (Prism 7) with individual biological replicates are shown and the mean is plotted as a bar. For statistical analysis, Student’s two-tailed t-test was used. P-values below 0.05 (p<0.05) were considered significant. All statistical analyses were performed in GraphPad Prism.

### Poly(A) enrichment

Poly(A) enrichment was performed using NEBNext^®^ Poly(A) mRNA Magnetic Isolation Module (New England BioLabs^®^ Inc.) according to manufacturer’s protocol with five μg total RNA from CTRL-KD or SFPQ-KD in HEK293T cells used as input.

### RNA sequencing

For total RNA sequencing of HEK293T, RNA from SFPQ, NONO, and CTRL KD (using two different siRNAs for each condition with biological duplicates) were rRNA depleted using RiboCop rRNA Depletion Kit V1.2 (Lexogen) according to manufacturer’s protocol. Subsequent cDNA libraries were prepared using SENSE Total RNA-Seq Library Prep Kit (Lexogen) following manufacturer’s protocol.

For 3’end sequencing, cDNA libraries from HEK293T were prepared using QuantSeq 3’ mRNA-Seq Library Prep Kit (Lexogen). For both methods, RNA quality was determined using the BioAnalyzer RNA nanochip (Agilent) and library concentration was quantified with KAPA Library Quant KIT RT-qPCR (Roche). Total RNAseq was done as 100nt paired-end sequencing and performed using the Illumnia platform (HiSEQ4000, BGI, Copenhagen), while for 3’end sequencing, 75nt single-end sequencing was performed at MOMA (Aarhus University Hospital) on a NextSeq500. For total RNA sequencing of HepG2 cells, library preparation and sequencing was performed at BGI (Copenhagen) using BGIseq.

### RNA-immunoprecipitation (RIP)

RIP was performed as previously described (Rinn et al. 2007) with some modifications to immunoprecipitate endogenous SFPQ and NONO. HepG2 and HEK293T cells were grown to confluence in 15cm^2^ dishes. Cells were harvested by trypsinization and resuspended in 2ml PBS, 2ml nuclear isolation buffer (1.28 M sucrose; 40 mM Tris-HCl Ph 7.5; 20 mM MgCl_2_; 4% Triton X-100) and 6ml water on ice for 20 min (with frequent mixing). Nuclei were pelleted by centrifugation at 2,500G for 15 min. Nuclear pellet was resuspended in 1ml RIP buffer (150 mM KCl, 25 mM Tris-HCl, pH 7.4, 5 mM EDTA, 0.5% Triton X-100 and 5 mM dithiothreitol (DTT) supplemented with Ribolock (Thermo Fisher Scientific) and proteinase inhibitor cocktail (Roche). Resuspended nuclei were split into two fractions of 500 μl each (for Mock and IP). Nuclear membrane and debris were pelleted by centrifugation at 13,000 RPM for 20 min. Antibody to SFPQ (P2860 Sigma), NONO (ab70335 Abcam) or FLAG epitope (Mock IP, F1804 Sigma) was added to supernatant (2.5μg) and incubated for 4hrs at 4C with gentle rotation. 20 μl of protein A/G beads were added and incubated for 1hr at 4C with gentle rotation. Beads isolated using magnetic, the supernatant were removed and beads were resuspended in 500μl RIP buffer and repeated for a total of 5 RIP washes. Beads were divided into two fractions for protein (30%) and RNA (70%). Protein fraction was resuspended in 2xSDS loading buffer [125 mM Tris–HCl pH 6.8, 20% glycerol, 5% SDS, and 0.2 M DTT] and boiled at 95°C for 5 min. RNA fraction was resuspended in 1ml TRIzol^®^ Reagent (Thermo Fisher Scientific).

### Western blotting

Cells were harvested in 1xPBS and centrifuged at 1200 rpm at 4°C for 5 min. For cell lysis, the cell pellet was collected and resuspended in 2xSDS loading buffer [125 mM Tris–HCl pH 6.8, 20% glycerol, 5% SDS, and 0.2 M DTT] and briefly boiled at 95°C for 5 min before loading 1% on a 10% Tris-Glycine SDS-PAGE gel (Thermo Fisher Scientific) and run for app. 1.5 hours at 125 V. The proteins were transferred to an Immobilon-P Transfer Membrane (EMD Millipore) by wet-blotting ON at 4°C at 25 V. Subsequently, the membrane was pre-blocked for 1 hr at RT with 10% skim milk, followed by 1 hr incubation with primary antibody (Supplementary Table 7) and 1 hr with secondary antibody. After each antibody incubation, the membrane was rinsed 3x 5 min in 1xPBS+0.05% Tween20 and 1x 5 min wash with 1xPBS. The protein bands were developed using SuperSignal West Femto Maximum Sensitivity Substrate kit (Thermo Fisher Scientific) and Amersham Hyperfilm ECL (GE Healthcare) or Medical film (MG-SR plus, Konica Minolta).

### Mapping, circRNA detection and quantification

Reads were mapped onto hg19 and mm10 for human and mouse data, respectively, with STAR (v2.7, (Dobin *et al*, 2013)) and quantified with featureCounts (v2.0.0, (Liao *et al*, 2014)) [featureCounts –p –O `–i gene_id –t exon`] using gencode annotations (v28lift37 for hg19 and v12 for mm10).

CircRNAs were predicted and quantified using ciri2 (Gao, Wang, & Zhao, 2015) and find_circ v1.2 (https://github.com/marvin-jens/find_circ) adhering to default settings, and only the shared predictions with ciri2 quantification were kept for analysis. circRNAs were annotated using *annotate_circ.py* (python scripts used are available at github/ncrnalab/pyutils). Flanking intron lengths were based on the mean total distance to the flanking exons based on gencode annotation (in case of multiple annotated flanking introns, the mean length was used), and IAE distance is the shortest possible Alu-mediated inverted repeat distance based on RepeatMasker (UCSC Genome Browser). For mouse, the IAE-distance is the shortest distance involving B1, B2 or B4 elements possible. DALI and PASI circRNAs were classified based on the median flanking intron lengths and median IAE distance in the sample. If no flanking introns were annotated, the circRNA was classified as ‘other’. Furthermore, circRNAs were classified as conserved if both splice sites coincide exactly with previously detected mouse circRNAs (Stagsted *et al*, 2019) converted to hg19 coordinates using the liftOver tool (UCSC genome browser). Flanking linear spliced reads from the circRNA producing loci were extracted using *get_flanking spliced_reads.py*.

### Cryptic/alternative splicing and intron retention

First, all spliced reads were extracted from bam-files using *get_spliced_reads.py* requiring at least an 8 nucleotide match on each exon. Then, separately for each splice-donor and –acceptor, all possible conjoining splice sites were extracted and counted using *get_alternative_splicing.py*. For each splice site, the most abundant splicing event across all samples was denoted as canonical, whereas all other splicing events from that particular splice site were either classified as ‘inclusion’ if shorter or ‘skipping’ if longer than the canonical.

Based on the output from alternative splicing, for each splice-site the intronic region of the shortest alternative event was quantified using featureCounts (as above) but with [`–minOverlap 5`] to avoid quantification of any overlapping spliced reads.

### eCLIP analysis

Pre-mapped eCLIP datasets were downloaded from encodeproject.org (hg19, see Supplementary Table 2). Based on the top1000 expressed circRNAs from HepG2 and K562 (See Supplementary Table 1), all reads aligning within 2000 nt upstream of the circRNA splice-acceptors or within 2000 nts downstream the circRNA splice-donors were counted using featureCounts. The same analysis was performed on all other annotated exons from the circRNAs host genes excluding the first and last host gene exons as well as exons involved in backsplicing. Moreover, genome-wide, exon-pairs were subsampled from gencode annotations to match the distributions of flanking intron lengths and linear spliced reads of DALI circRNAs; DALI-like exons. Enrichment was assessed by Wilcoxon rank-sum test between the number of reads flanking circRNAs compared to host exons. In case of SFPQ eCLIP, the two replicate samples were normalized to total number of aligned reads and then divided by the normalized input sample counts to obtain an IP/Input value for each locus.

### Quantseq analysis

Quantseq reads were mapped onto hg19 using STAR as described above. The resulting bam-files were merged and divided into plus and minus-strand alignments. Then, using MACS2 peakcall (v2.2.6, https://github.com/macs3-project/MACS) with parameters [`--nomodel --shift 0 –g 2.9e9`], quantseq peaks were extracted. Each peak was then subsequently quantified using featureCounts and analysed for the presence of polyA signal (A[AU]UAAA) and the presence of polyA-stretches within the locus or in the immediate flanking regions (+/− 50 nts). Based on gencode annotation each peak was assigned as exonic, intronic, ambiguous, or intergenic.

### Differential gene expression

First, in all analyses, low-expressed entries defined by mean counts across all samples <1 and expressed in less than three samples were discarded. Then, analysis of differential gene expression was performed using DESeq2 (v1.24.0, (Love *et al*, 2014)) using formula ~ treatment, where treatment denotes the knockdown/knockout target. For mRNA and circRNA expression, the raw counts were merged and analysed in bulk. For conditional analysis, such as circ vs linear, alternative vs canonical splicing, and intron-retention vs intron-splicing, raw counts for each type was combined in one expression matrix with the associated design formula: ~ treatment * type, where type denotes circular or linear splicing (in case of circ vs linear). The log2FoldChange and p-adjust values from the interaction-term (treatment:type) was used in subsequent analyses. For binned analysis of transcripts, each locus was sliced and re-annotated as 20 equally sized bins irrespective of exon-intron structure, and this was then used in the featureCounts quantification. After differential expression analysis by DESeq2, genes were subgrouped into five k-means clusters based on the DESeq2-derived log2foldchange of all 20 bins.

An adjusted p-value below 0.05 was considered significant. All statistics were conducted in R (v3.6.3) and visualizations were done in R using ggplot2 (v3.3.0) and GraphPad.

## Data Accessibility

Sequencing data will be uploaded to the GEO omnibus, and scripts for RNAseq data processing are available at github: github.com/ncrnalab/pyutils

## Acknowledgements

We would like to thank Dr. Anne Færch Nielsen, Dr. Karoline Kragh Ebbesen and Professor Jørgen Kjems for constructive comments and critical reading of the manuscript. This work was supported by the Novo Nordisk Foundation (NNF16OC0019874 to T.B.H) and the circRTrain ITN network.

## Author contributions

LVWS performed western blot and library preparation. ETL performed RIP analysis. LVWS and ETL performed knockdown and qRT-PCR. TBH performed RNAseq analyses and supervised the project. LVWS, ETL and TBH wrote the manuscript.

## Bibliography

Ajuh P, Kuster B, Panov K, Zomerdijk JC, Mann M & Lamond AI (2000) Functional analysis of the human CDC5L complex and identification of its components by mass spectrometry. EMBO J. 19: 6569–6581

Aktaş T, Avşar Iliki, Maticzka D, Bhardwaj V, Pessoa Rodrigues C, Mittler G, Manke T, Backofen R & Akhtar A (2017) DHX9 suppresses RNA processing defects originating from the Alu invasion of the human genome. Nature 544: 115–119

Ashwal-Fluss R, Meyer M, Pamudurti NR, Ivanov A, Bartok O, Hanan M, Evantal N, Memczak S, Rajewsky N & Kadener S (2014) circRNA biogenesis competes with pre-mRNA splicing. Mol. Cell 56: 55–66

Buxadé M, Morrice N, Krebs DL & Proud CG (2008) The PSF.p54nrb complex is a novel Mnk substrate that binds the mRNA for tumor necrosis factor alpha. J. Biol. Chem. 283: 57–65

Clemson CM, Hutchinson JN, Sara SA, Ensminger AW, Fox AH, Chess A & Lawrence JB (2009) An architectural role for a nuclear noncoding RNA: NEAT1 RNA is essential for the structure of paraspeckles. Mol. Cell 33: 717–726

Conn SJ, Pillman KA, Toubia J, Conn VM, Salmanidis M, Phillips CA, Roslan S, Schreiber AW, Gregory PA & Goodall GJ (2015) The RNA binding protein quaking regulates formation of circRNAs. Cell 160: 1125–1134

Dobin A, Davis CA, Schlesinger F, Drenkow J, Zaleski C, Jha S, Batut P, Chaisson M & Gingeras TR (2013) STAR: ultrafast universal RNA-seq aligner. Bioinformatics 29: 15–21

Dong B, Horowitz DS, Kobayashi R & Krainer AR (1993) Purification and cDNA cloning of HeLa cell p54nrb, a nuclear protein with two RNA recognition motifs and extensive *homology to human splicing factor PSF and Drosophila NONA/BJ6*. Nucleic Acids Res. 21: 4085–4092

Dubin RA, Kazmi MA & Ostrer H (1995) Inverted repeats are necessary for circularization of the mouse testis Sry transcript. Gene 167: 245–248

Emili A, Shales M, McCracken S, Xie W, Tucker PW, Kobayashi R, Blencowe BJ & Ingles CJ (2002) Splicing and transcription-associated proteins PSF and p54nrb/nonO bind to the RNA polymerase II CTD. RNA 8: 1102–1111

ENCODE Project Consortium (2012) An integrated encyclopedia of DNA elements in the human genome. Nature 489: 57–74

Errichelli L, Dini Modigliani S, Laneve P, Colantoni A, Legnini I, Capauto D, Rosa A, De Santis R, Scarfò R, Peruzzi G, Lu L, Caffarelli E, Shneider NA, Morlando M & Bozzoni I (2017) FUS affects circular RNA expression in murine embryonic stem cell-derived motor neurons. Nat. Commun. 8: 14741

Fox AH, Nakagawa S, Hirose T & Bond CS (2018) Paraspeckles: where long noncoding RNA meets phase separation. Trends Biochem. Sci. 43: 124–135

Furukawa MT, Sakamoto H & Inoue K (2015) Interaction and colocalization of HERMES/RBPMS with NonO, PSF, and G3BP1 in neuronal cytoplasmic RNP granules in mouse retinal line cells. Genes Cells 20: 257–266

Gozani O, Patton JG & Reed R (1994) A novel set of spliceosome-associated proteins and the essential splicing factor PSF bind stably to pre-mRNA prior to catalytic step II of the splicing reaction. EMBO J. 13: 3356–3367

Hansen TB, Jensen TI, Clausen BH, Bramsen JB, Finsen B, Damgaard CK & Kjems J (2013) Natural RNA circles function as efficient microRNA sponges. Nature 495: 384–388

Henriques T, Gilchrist DA, Nechaev S, Bern M, Muse GW, Burkholder A, Fargo DC & Adelman K (2013) Stable pausing by RNA polymerase II provides an opportunity to target and integrate regulatory signals. Mol. Cell 52: 517–528

Hosokawa M, Takeuchi A, Tanihata J, Iida K, Takeda S & Hagiwara M (2019) Loss of RNA-Binding Protein Sfpq Causes Long-Gene Transcriptopathy in Skeletal Muscle and Severe Muscle Mass Reduction with Metabolic Myopathy. iScience 13: 229–242

Iida K, Hagiwara M & Takeuchi A (2020) Multilateral Bioinformatics Analyses Reveal the Function-Oriented Target Specificities and Recognition of the RNA-Binding Protein SFPQ. iScience 23: 101325

Ishigaki S, Fujioka Y, Okada Y, Riku Y, Udagawa T, Honda D, Yokoi S, Endo K, Ikenaka K, Takagi S, Iguchi Y, Sahara N, Takashima A, Okano H, Yoshida M, Warita H, Aoki M, Watanabe H, Okado H, Katsuno M, et al (2017) Altered Tau Isoform Ratio Caused by Loss of FUS and SFPQ Function Leads to FTLD-like Phenotypes. Cell Rep. 18: 1118–1131

Ito T, Watanabe H, Yamamichi N, Kondo S, Tando T, Haraguchi T, Mizutani T, Sakurai K, Fujita S, Izumi T, Isobe T & Iba H (2008) Brm transactivates the telomerase reverse transcriptase (TERT) gene and modulates the splicing patterns of its transcripts in concert with p54(nrb). Biochem. J. 411: 201–209

Ivanov A, Memczak S, Wyler E, Torti F, Porath HT, Orejuela MR, Piechotta M, Levanon EY, Landthaler M, Dieterich C & Rajewsky N (2015) Analysis of intron sequences reveals hallmarks of circular RNA biogenesis in animals. Cell Rep. 10: 170–177

Jeck WR, Sorrentino JA, Wang K, Slevin MK, Burd CE, Liu J, Marzluff WF & Sharpless NE (2013) Circular RNAs are abundant, conserved, and associated with ALU repeats. RNA 19:141–157

Kameoka S, Duque P & Konarska MM (2004) p54(nrb) associates with the 5’ splice site within large transcription/splicing complexes. EMBO J. 23: 1782–1791

Kaneko S, Rozenblatt-Rosen O, Meyerson M & Manley JL (2007) The multifunctional protein p54nrb/PSF recruits the exonuclease XRN2 to facilitate pre-mRNA 3’ processing and transcription termination. Genes Dev. 21: 1779–1789

Knott GJ, Bond CS & Fox AH (2016) The DBHS proteins SFPQ, NONO and PSPC1: a multipurpose molecular scaffold. Nucleic Acids Res. 44: 3989–4004

Knott GJ, Lee M, Passon DM, Fox AH & Bond CS (2015) Caenorhabditis elegans NONO-1: Insights into DBHS protein structure, architecture, and function. Protein Sci. 24: 2033–2043

Kramer MC, Liang D, Tatomer DC, Gold B, March ZM, Cherry S & Wilusz JE (2015) Combinatorial control of Drosophila circular RNA expression by intronic repeats, hnRNPs, and SR proteins. Genes Dev. 29: 2168–2182

Kwiatkowski TJ, Bosco DA, Leclerc AL, Tamrazian E, Vanderburg CR, Russ C, Davis A, Gilchrist J, Kasarskis EJ, Munsat T, Valdmanis P, Rouleau GA, Hosler BA, Cortelli P, de Jong PJ, Yoshinaga Y, Haines JL, Pericak-Vance MA, Yan J, Ticozzi N, et al (2009) Mutations in the FUS/TLS gene on chromosome 16 cause familial amyotrophic lateral sclerosis. Science 323: 1205–1208

Lagier-Tourenne C, Polymenidou M, Hutt KR, Vu AQ, Baughn M, Huelga SC, Clutario KM, Ling S-C, Liang TY, Mazur C, Wancewicz E, Kim AS, Watt A, Freier S, Hicks GG, Donohue JP, Shiue L, Bennett CF, Ravits J, Cleveland DW, et al (2012) Divergent roles of ALS-linked proteins FUS/TLS and TDP-43 intersect in processing long pre-mRNAs. Nat. Neurosci. 15: 1488–1497

Lee M, Sadowska A, Bekere I, Ho D, Gully BS, Lu Y, Iyer KS, Trewhella J, Fox AH & Bond CS (2015) The structure of human SFPQ reveals a coiled-coil mediated polymer essential for functional aggregation in gene regulation. Nucleic Acids Res. 43: 3826–3840

Liang D, Tatomer DC, Luo Z, Wu H, Yang L, Chen L-L, Cherry S & Wilusz JE (2017) The Output of Protein-Coding Genes Shifts to Circular RNAs When the Pre-mRNA Processing Machinery Is Limiting. Mol. Cell 68: 940–954.e3

Liao Y, Smyth GK & Shi W (2014) featureCounts: an efficient general purpose program for assigning sequence reads to genomic features. Bioinformatics 30: 923–930

Love MI, Huber W & Anders S (2014) Moderated estimation of fold change and dispersion for RNA-seq data with DESeq2. Genome Biol. 15: 550

Luisier R, Tyzack GE, Hall CE, Mitchell JS, Devine H, Taha DM, Malik B, Meyer I, Greensmith L, Newcombe J, Ule J, Luscombe NM & Patani R (2018) Intron retention and nuclear loss of SFPQ are molecular hallmarks of ALS. Nat. Commun. 9: 2010

Makarov EM, Makarova OV, Urlaub H, Gentzel M, Will CL, Wilm M & Lührmann R (2002) Small nuclear ribonucleoprotein remodeling during catalytic activation of the spliceosome. Science 298: 2205–2208

Memczak S, Jens M, Elefsinioti A, Torti F, Krueger J, Rybak A, Maier L, Mackowiak SD, Gregersen LH, Munschauer M, Loewer A, Ziebold U, Landthaler M, Kocks C, le Noble F & Rajewsky N (2013) Circular RNAs are a large class of animal RNAs with regulatory potency. Nature 495: 333–338

Mora Gallardo C, Sánchez de Diego A, Gutiérrez Hernández J, Talavera-Gutiérrez A, Fischer T, Martínez-A C & van Wely KHM (2019) Dido3-dependent SFPQ recruitment maintains efficiency in mammalian alternative splicing. Nucleic Acids Res. 47: 5381–5394

Nakaya T, Alexiou P, Maragkakis M, Chang A & Mourelatos Z (2013) FUS regulates genes coding for RNA-binding proteins in neurons by binding to their highly conserved introns. RNA 19: 498–509

Palangat M, Hittinger CT & Landick R (2004) Downstream DNA selectively affects a paused conformation of human RNA polymerase II. J. Mol. Biol. 341: 429–442

Passon DM, Lee M, Rackham O, Stanley WA, Sadowska A, Filipovska A, Fox AH & Bond CS (2012) Structure of the heterodimer of human NONO and paraspeckle protein *component 1 and analysis of its role in subnuclear body formation*. Proc Natl Acad Sci USA 109:4846–4850

Patton JG, Porro EB, Galceran J, Tempst P & Nadal-Ginard B (1993) Cloning and characterization of PSF, a novel pre-mRNA splicing factor. Genes Dev. 7: 393–406

Peng R, Dye BT, Pérez I, Barnard DC, Thompson AB & Patton JG (2002) PSF and p54nrb bind a conserved stem in U5 snRNA. RNA 8: 1334–1347

Ragan C, Goodall GJ, Shirokikh NE & Preiss T (2019) Insights into the biogenesis and potential functions of exonic circular RNA. Sci. Rep. 9: 2048

Rosonina E, Ip JYY, Calarco JA, Bakowski MA, Emili A, McCracken S, Tucker P, Ingles CJ & Blencowe BJ (2005) Role for PSF in mediating transcriptional activator-dependent stimulation of pre-mRNA processing in vivo. Mol. Cell. Biol. 25: 6734–6746

Rybak-Wolf A, Stottmeister C, Glažar P, Jens M, Pino N, Giusti S, Hanan M, Behm M, Bartok O, Ashwal-Fluss R, Herzog M, Schreyer L, Papavasileiou P, Ivanov A, Öhman M, Refojo D, Kadener S & Rajewsky N (2015) Circular RNAs in the mammalian brain are highly abundant, conserved, and dynamically expressed. Mol. Cell 58: 870–885

Salzman J, Chen RE, Olsen MN, Wang PL & Brown PO (2013) Cell-type specific features of circular RNA expression. PLoS Genet. 9: e1003777

Salzman J, Gawad C, Wang PL, Lacayo N & Brown PO (2012) Circular RNAs are the predominant transcript isoform from hundreds of human genes in diverse cell types. PLoS ONE 7: e30733

Stagsted LV, Nielsen KM, Daugaard I & Hansen TB (2019) Noncoding AUG circRNAs constitute an abundant and conserved subclass of circles. Life Sci. Alliance 2:

Szabo L, Morey R, Palpant NJ, Wang PL, Afari N, Jiang C, Parast MM, Murry CE, Laurent LC & Salzman J (2015) Statistically based splicing detection reveals neural enrichment and tissue-specific induction of circular RNA during human fetal development. Genome Biol. 16: 126

Takeuchi A, Iida K, Tsubota T, Hosokawa M, Denawa M, Brown JB, Ninomiya K, Ito M, Kimura H, Abe T, Kiyonari H, Ohno K & Hagiwara M (2018) Loss of Sfpq Causes Long-Gene Transcriptopathy in the Brain. Cell Rep. 23: 1326–1341

Thomas-Jinu S, Gordon PM, Fielding T, Taylor R, Smith BN, Snowden V, Blanc E, Vance C, Topp S, Wong C-H, Bielen H, Williams KL, McCann EP, Nicholson GA, Pan-Vazquez A, Fox AH, Bond CS, Talbot WS, Blair IP, Shaw CE, et al (2017) Non-nuclear Pool of Splicing Factor SFPQ Regulates Axonal Transcripts Required for Normal Motor Development. Neuron 94: 931

Urban RJ, Bodenburg Y, Kurosky A, Wood TG & Gasic S (2000) Polypyrimidine tract-binding protein-associated splicing factor is a negative regulator of transcriptional activity of the porcine p450scc insulin-like growth factor response element. Mol. Endocrinol. 14: 774–782

Vance C, Rogelj B, Hortobágyi T, De Vos KJ, Nishimura AL, Sreedharan J, Hu X, Smith B, Ruddy D, Wright P, Ganesalingam J, Williams KL, Tripathi V, Al-Saraj S, Al-Chalabi A, Leigh PN, Blair IP, Nicholson G, de Belleroche J, Gallo J-M, et al (2009) Mutations in FUS, an RNA processing protein, cause familial amyotrophic lateral sclerosis type 6. Science 323: 1208–1211

Venø MT, Hansen TB, Venø ST, Clausen BH, Grebing M, Finsen B, Holm IE & Kjems J (2015) Spatio-temporal regulation of circular RNA expression during porcine embryonic brain development. Genome Biol. 16: 245

Westholm JO, Miura P, Olson S, Shenker S, Joseph B, Sanfilippo P, Celniker SE, Graveley BR & Lai EC (2014) Genome-wide analysis of drosophila circular RNAs reveals their structural and sequence properties and age-dependent neural accumulation. Cell Rep. 9: 1966–1980

You X, Vlatkovic I, Babic A, Will T, Epstein I, Tushev G, Akbalik G, Wang M, Glock C, Quedenau C, Wang X, Hou J, Liu H, Sun W, Sambandan S, Chen T, Schuman EM & Chen W (2015) Neural circular RNAs are derived from synaptic genes and regulated by development and plasticity. Nat. Neurosci. 18: 603–610

Zhang X-O, Wang H-B, Zhang Y, Lu X, Chen L-L & Yang L (2014) Complementary sequence-mediated exon circularization. Cell 159: 134–147

Zhang Y, Xue W, Li X, Zhang J, Chen S, Zhang J-L, Yang L & Chen L-L (2016) The biogenesis of nascent circular rnas. Cell Rep. 15: 611–624

Zhang Z & Carmichael GG (2001) The fate of dsRNA in the nucleus: a p54(nrb)-containing complex mediates the nuclear retention of promiscuously A-to-I edited RNAs. Cell 106: 465–475

