## Supplementary Figures 1-14 for "The RNA-binding protein SFPQ preserves long-intron splicing and regulates circRNA biogenesis"

Supplementary Figure 1

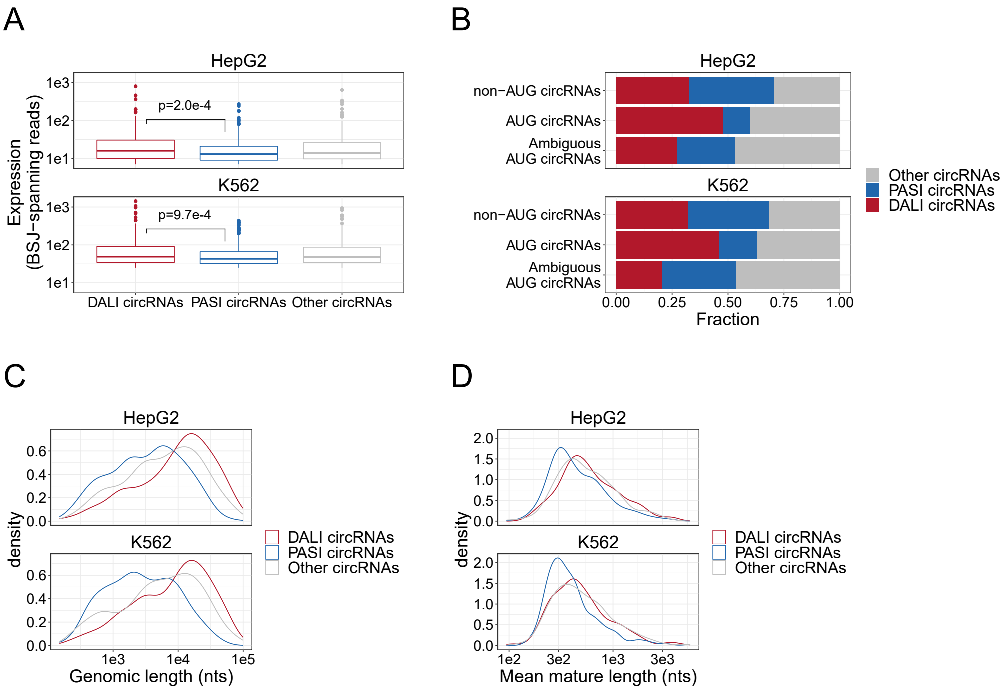


**Supplementary Figure 1: circRNAome in HepG2 and K562 from ENCODE. A)** Boxplot showing expression distribution of top1000 expressed circRNA as measured by back-splice junction (BSJ) spanning reads for DALI, PASI and Other circRNAs in HepG2 and K562 cells. **B)** The fraction of DALI, PASI and Other circRNAs comprising the previously characterized subset of conserved circRNAs, the AUG circRNAs (Stagsted et al, 2019). **C-D)** The distribution of genomic lengths, i.e. the genomic distance between the SD and SA involved in backsplicing (C) and the mature length, i.e. the predicted length of the fully spliced circRNAs (D) stratified by subgroup as denoted.

Supplementary Figure 2

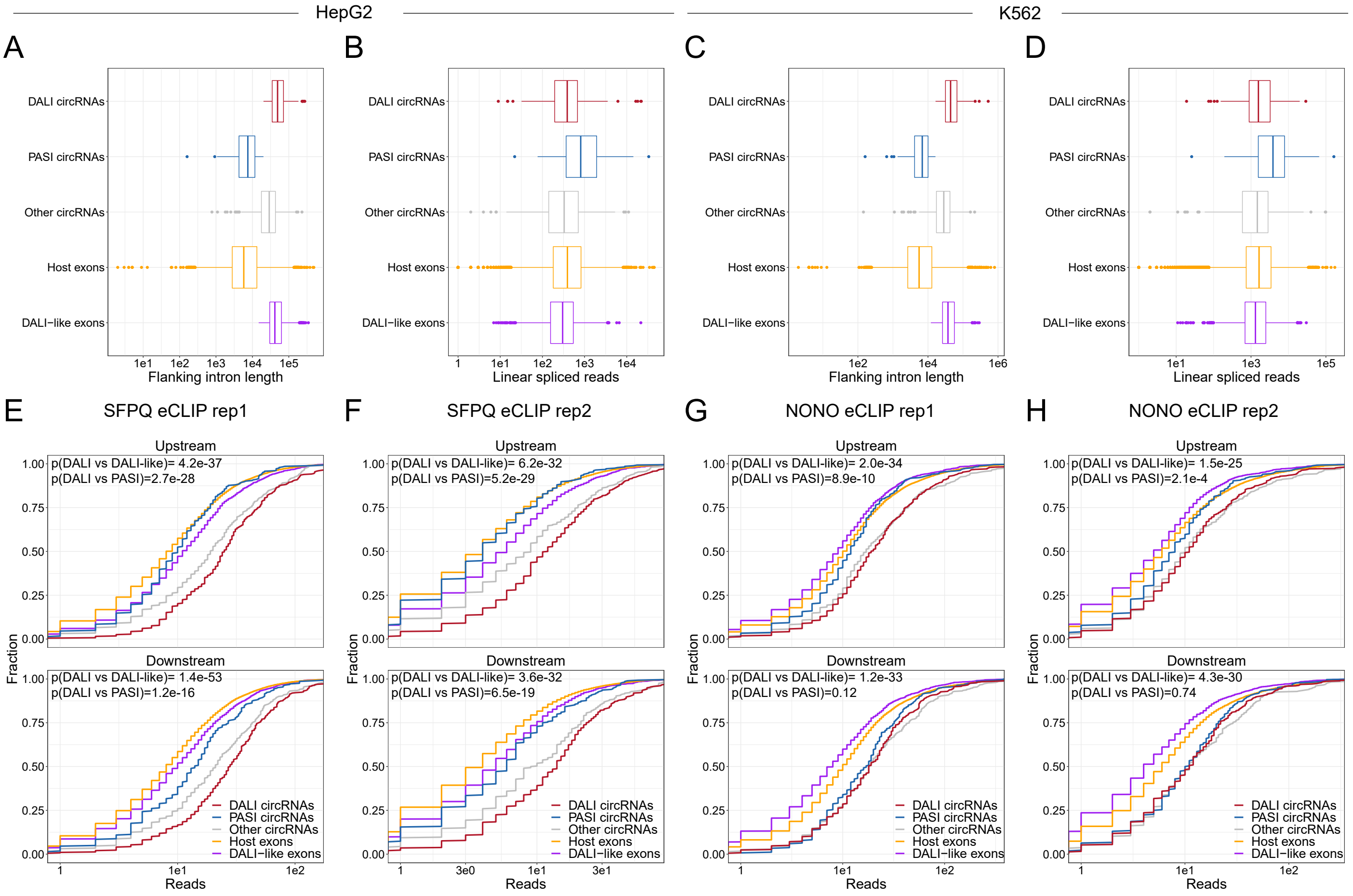


**Supplementary Figure 2: SFPQ and NONO enriched on circRNA flanking introns. A-D)** For HepG2 (A and B) and K562 (C and D), boxplots showing the distribution of flanking intron length (A and C) or linear spliced reads (B and D) for DALI circRNAs (red), PASI circRNAs (blue), other circRNAs (grey), host exons, i.e. all other annotated exons from the circRNA-producing loci (orange), and DALI-like circRNAs, i.e. exon-pairs from annotated genes sampled to resemble DALI circRNAs based on flanking intron lengths and linear spliced reads (purple). **E-H)** Boxplots of reads from SFPQ eCLIP rep1 (F), SFPQ eCLIP rep2 (G), NONO eCLIP rep1 (H), and NONO eCLIP rep2 associated with each subgroup in HepG2 cells (F-G) and K562 cells (H-I) stratified by upstream (upper facets) and downstream (lower facets) aligned reads. P-values are calculated using Wilcoxon rank-sum tests.

Supplementary Figure 3

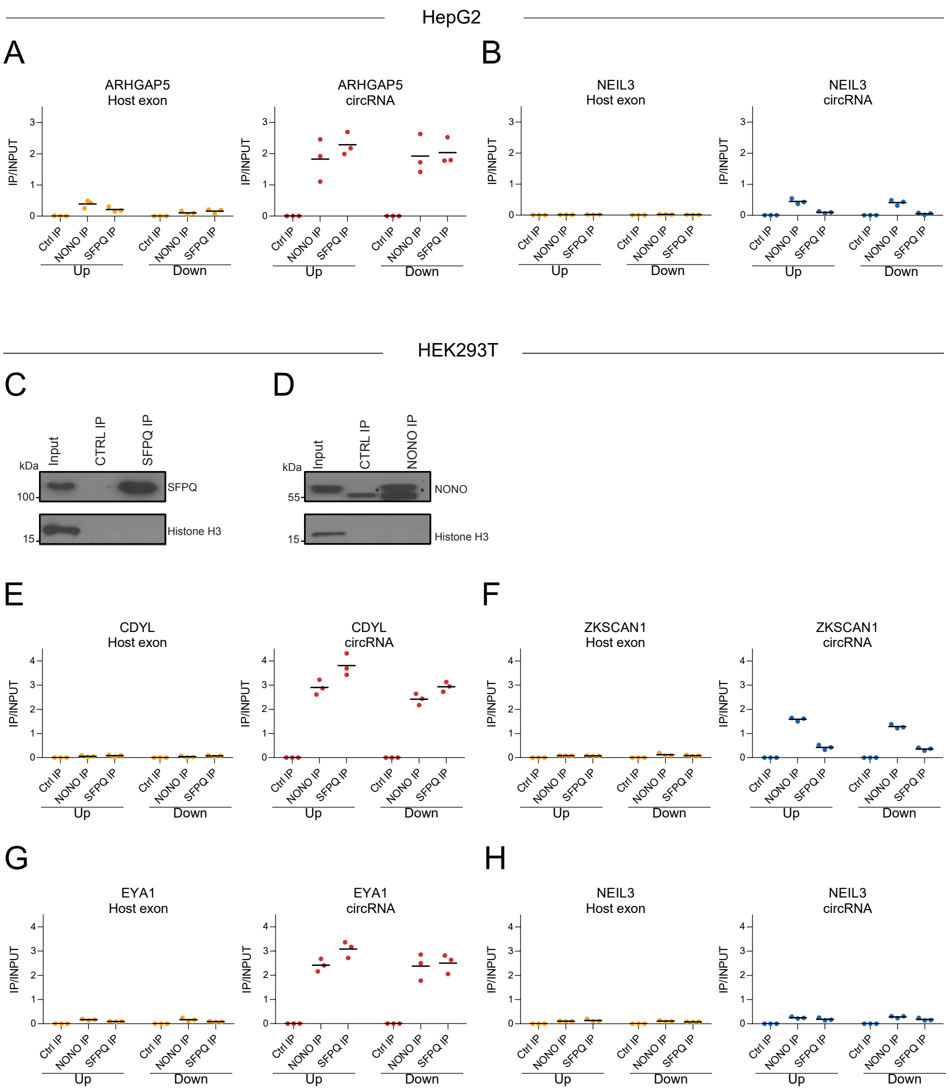


**Supplementary Figure 3: RNA immunoprecipitation of SFPQ and NONO confirms enrichment. A-B)** As in Fig. 1G-H, qRT-PCR on denoted intronic regions in ARHGAP5 (A) and NEIL3 (B) transcripts upon RNA IP of endogenous SFPQ or NONO from nuclear fractions of HepG2 cells. **C-D)** Western blotting of endogenous immunoprecipitated (IP) SFPQ (C) or NONO (D) from nuclear fractions of HEK293T cells with Histone H3 as a loading control. Asterisks denote bands derived from the IP antibody. **E-H)** As in A-B but using HEK293T cells and with qRT-PCR on CDYL (C), ZKSCAN1 (D), EYA (E) and NEIL3 (F). Data for three biological replicates are shown.

Supplementary Figure 4

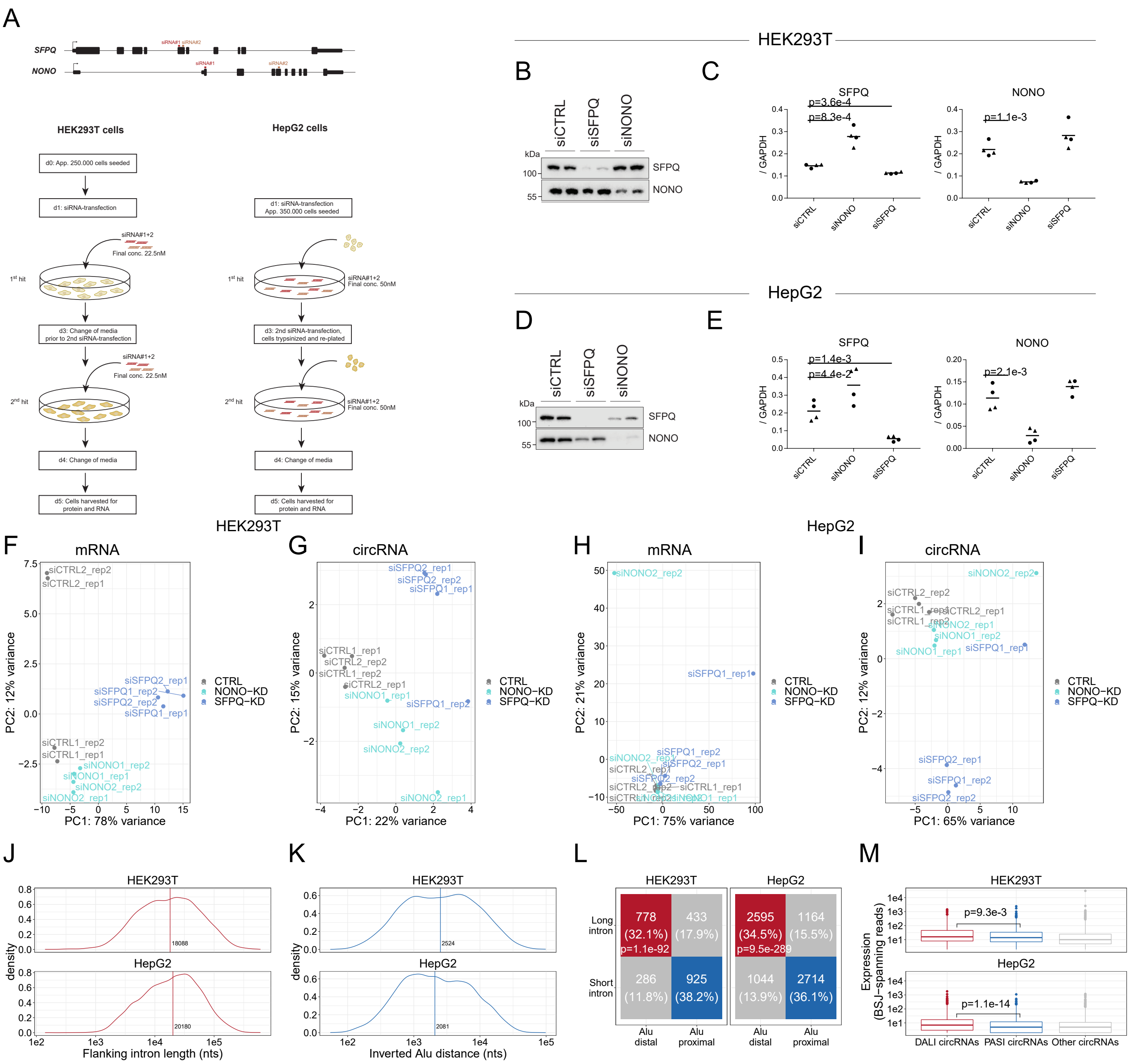


**Supplementary Figure 4: SFPQ/NONO-depletion in HEK293T and HepG2 cells. A)** Schematic showing the siRNA-knockdown protocol in HEK293T and HepG2 cells. For each condition (CTRL, NONO-KD and SFPQ-KD) two different siRNA designs were used to reduce off-targeting effects, and for each siRNA, the experiment was performed in biological duplicates. **B-E**) Western blotting (B and D) and qRT-PCR (C and E) validation of knockdown (2$^{nd}$ replicate) in HEK293T (B-C) and HepG2 (D-E) cells. Data for four biological replicates are shown. P-values are calculated using student's two-tailed t-test. **F-I)** PCA analysis of top500 most variable mRNAs (F and H) and circRNAs (G-I) as measured across samples in HEK293T cells (F and G) and HepG2 cell (H and I) subjected to SFPQ and NONO-depletion. The individual samples are color-coded by the knockdown target as denoted. **J-K)** Distributions of flanking intron lengths (J) and inverted *Alu* distances (K) for circRNAs detected in HEK293T (upper facet) and HepG2 (lower facet) cells. The vertical line and the corresponding value represents the median. **L)** Contingency table for circRNAs stratified by flanking intron lengths and inverted *Alu* distances in HEK293T (left facet) and HepG2 (right facet) cells. The table is color-coded by circRNA subgroups; DALI (red), PASI (blue) and the others (grey). **M)** Boxplot showing the number of BSJ-spanning reads for the top1000 circRNAs stratified by subgroup as denoted. P-values are calculated using Wilcoxon rank-sum tests.

Supplementary Figure 5

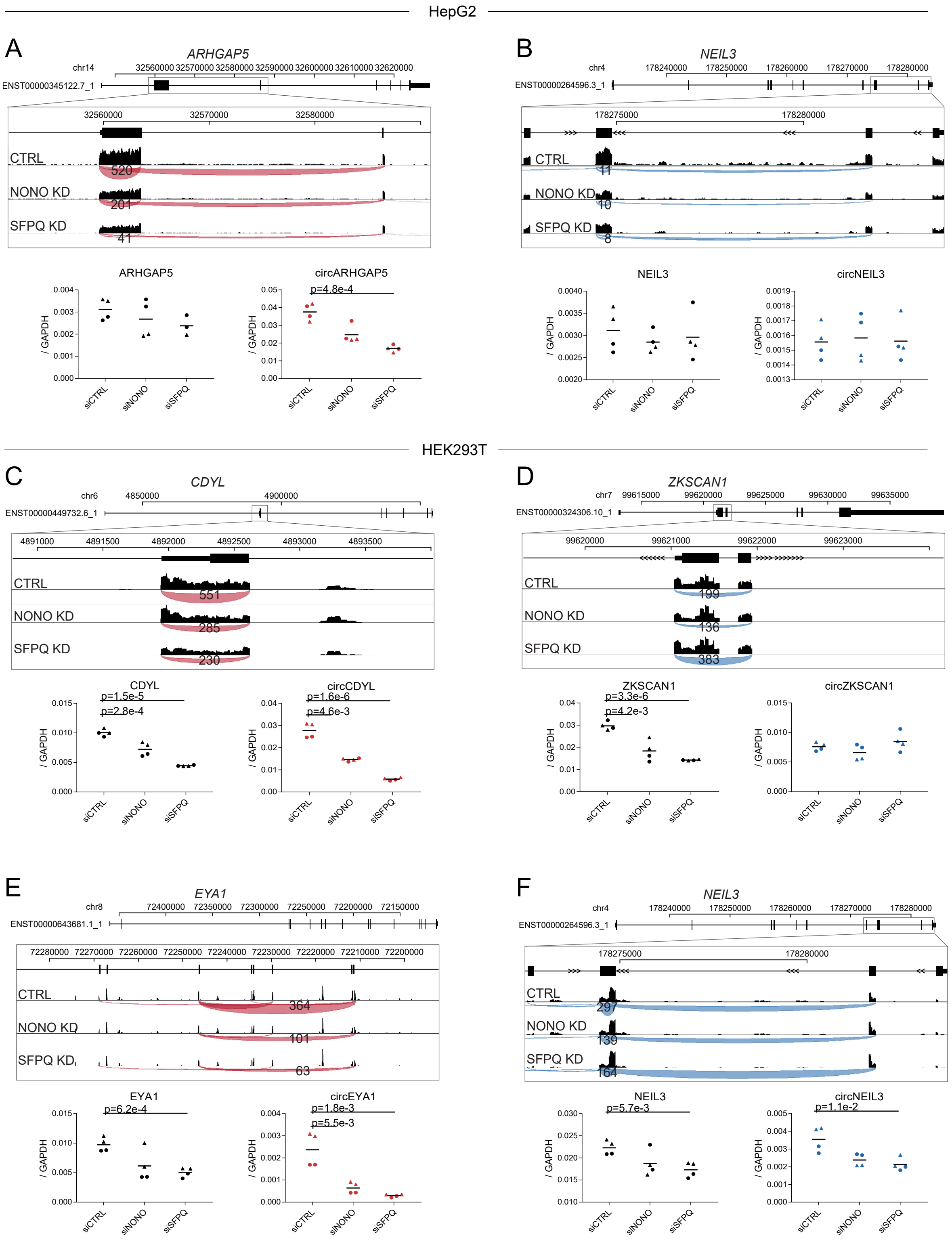


**Supplementary Figure 5: Expression profiles for selected circRNAs. A-F)** Genomic exon-intron structures of selected circRNAs-producing genes with screendumps showing circRNAs backsplicing reads obtained from RNAseq and visualized using IGV genome browser from HepG2 (A-B) and HEK293T (C-F). Below, qRT-PCR validation in independent experiment using BSJ-spanning primers (circRNAs expression) and flanking linear-splicing primers (host-gene expression) relative to *GAPDH.* Data for four biological replicates are shown. P-values are calculated using student's two-tailed t-test.

Supplementary Figure 6

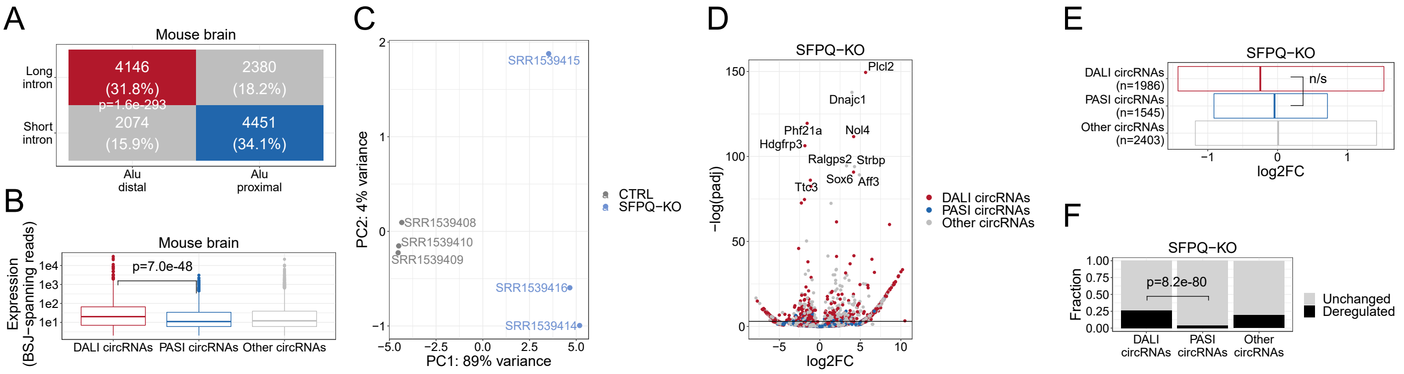


**Supplementary Figure 6: CircRNAome analysis of SFPQ knockout mouse brain data (GSE60246). A)** Two-by-two contingency table of circRNAs stratified by intron length and inverted *Alu* distance. P-value calculated by Fisher's exact test. **B)** Boxplot on the distribution of BSJ-spanning reads for each circRNAs subgroup. P-value calculated using Wilcoxon rank-sum test. **C)** PCA analysis of wild-type (CTRL) and SFPQ knockout (SFPQ-KO) samples based on circRNA expression. **D)** Volcano plot showing deregulated circRNA expression comparing WT (CTRL) and SFPQ-KO mouse color-coded by circRNA subgroup as denoted. **E)** Quantile plot showing .25, .5 (median) and .75 quantiles of the log2foldchange distribution between WT and SFPQ-KO for each circRNA subgroup. **F)** Barplot showing the fraction of circRNAs from each subgroup showing significant deregulation upon SFPQ knockout. P-value calculated using the Fisher's exact test.

Supplementary Figure 7

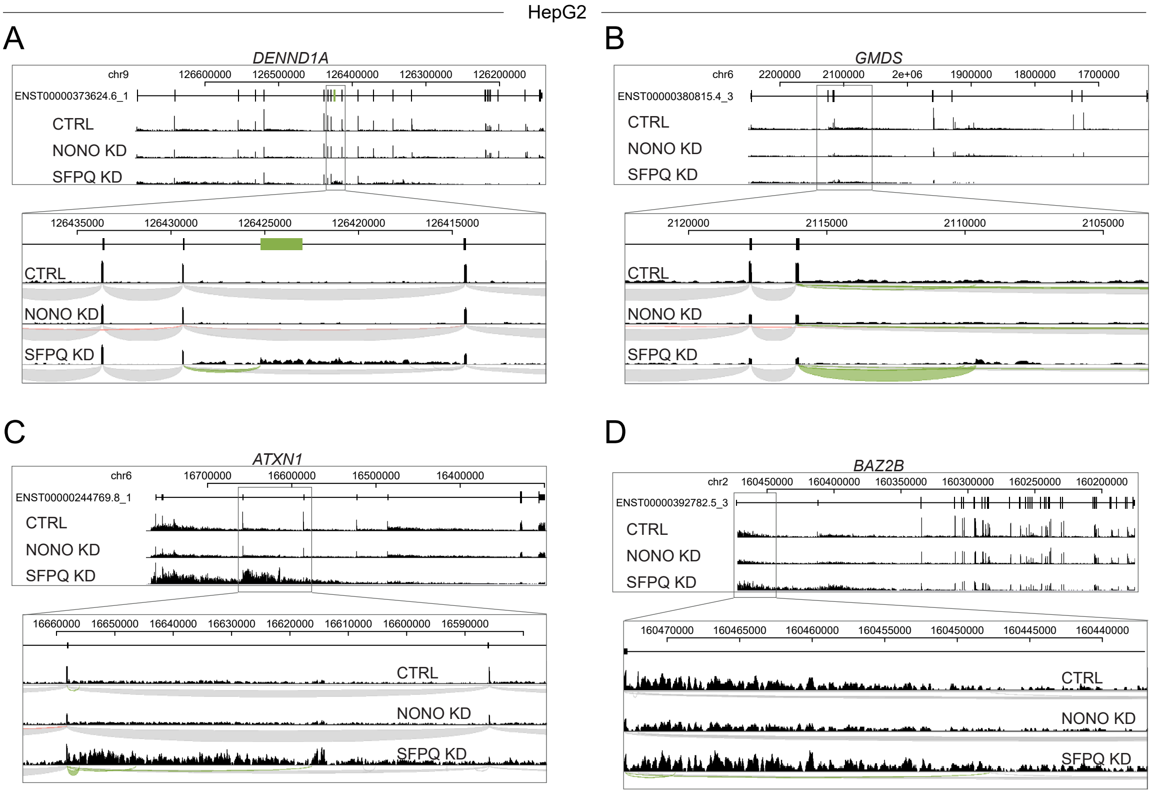


**Supplementary Figure 7: Genic expression profile for selected long genes. A-D)** Read coverage from HepG2 cells with either NONO- or SFPQ-depletion on DENND1A (A), GMDS (B), ATXN1 (C), and BAZ2B (D)**.** The tracks are composed of merged and normalized expression from all replicates/siRNA-designs.

### Supplementary Figure 8

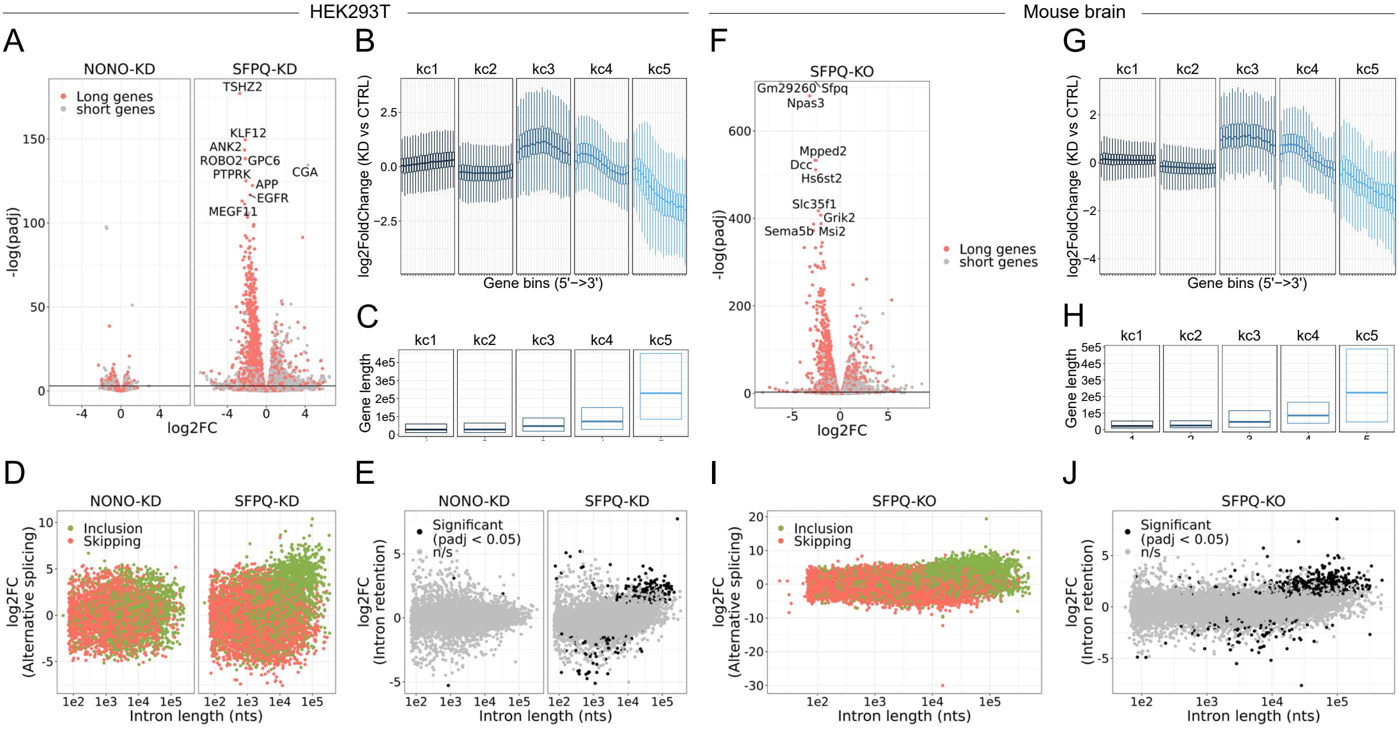


**Supplementary Figure 8: SFPQ ensures long-gene expression (HEK293T + MOUSE). A)** Volcano plot stratified by genes higher or lower than median gene length, where length is the annotated distance from promoter to terminator. **B)** Boxplot showing binned expression of clustered genes in SFPQ-depleted sampled relative to CTRL. Each gene is sliced into 20 bins, and the differential expression of each bin is determined and subgrouped into five kmeans clusters (see Methods). **C)** Boxplot showing gene lengths distribution stratified by clusters obtained in B. **D)** Scatter plot showing alternative splicing in NONO and SFPQ-depleted samples as a function of canonical intron length and color-coded by type of splicing (either inclusion or skipping, see schematic in Figure 4H)**. E)** Scatter plot showing effects on retention upon SFPQ and NONO depletion as a function of intron length. **F-J)** analyses as in A-E on mouse brain SFPQ knockout samples (GSE60246, see Supplementary Table 5).

Supplementary Figure 9

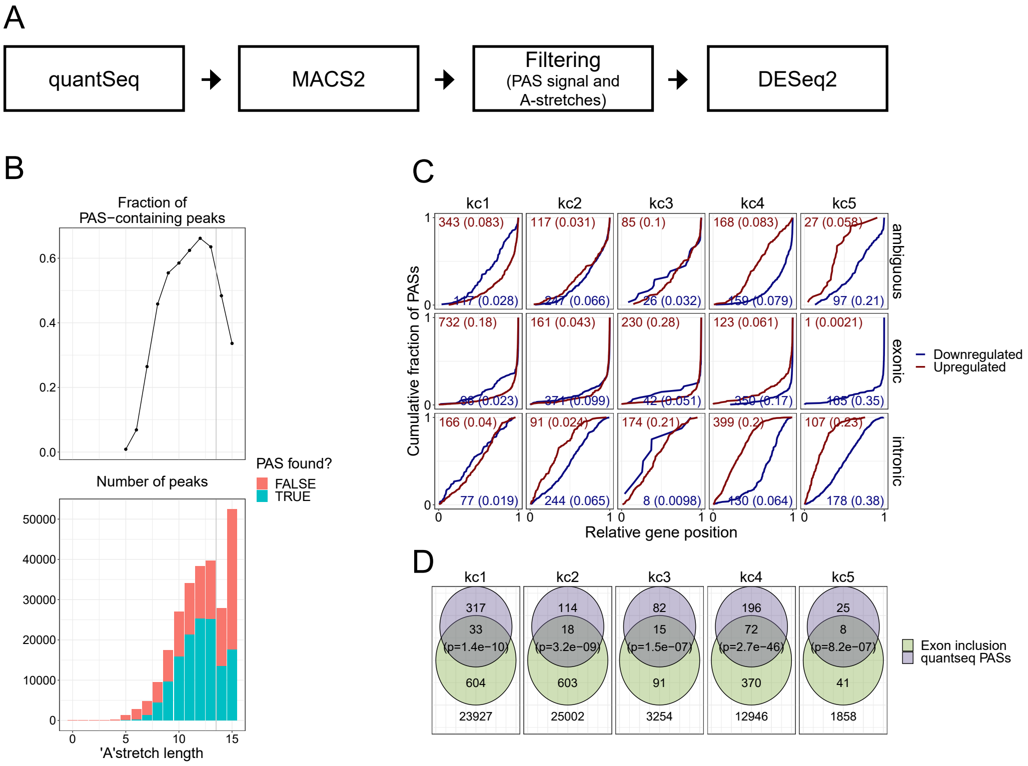


**Supplementary Figure 9: Quantseq analysis. A)** Schematic depicting the quantseq workflow **B)** Top: characterization of the fraction of PAS-containing peaks, where PAS is defined as AAUAAA or AUUAAA, as a function of longest oligo-A stretch identified in peak +/- 50 nt flanking region. Bottom: Total number of peaks identified with (green) or without (orange) PAS as a function of longest A-stretch. **C)** Venn diagrams (as in Fig. 5E) showing overlapping quantseq PASs and cryptic splicing but stratified into the five kmeans clusters. **D)** Relative quantseq PAS position within annotated genes (as in Fig. 5B) but stratified by kmeans clusters. Numbers denote the number of peaks in each group and the fraction of genes with significant de-regulated peaks in parenthesis.

Supplementary Figure 10

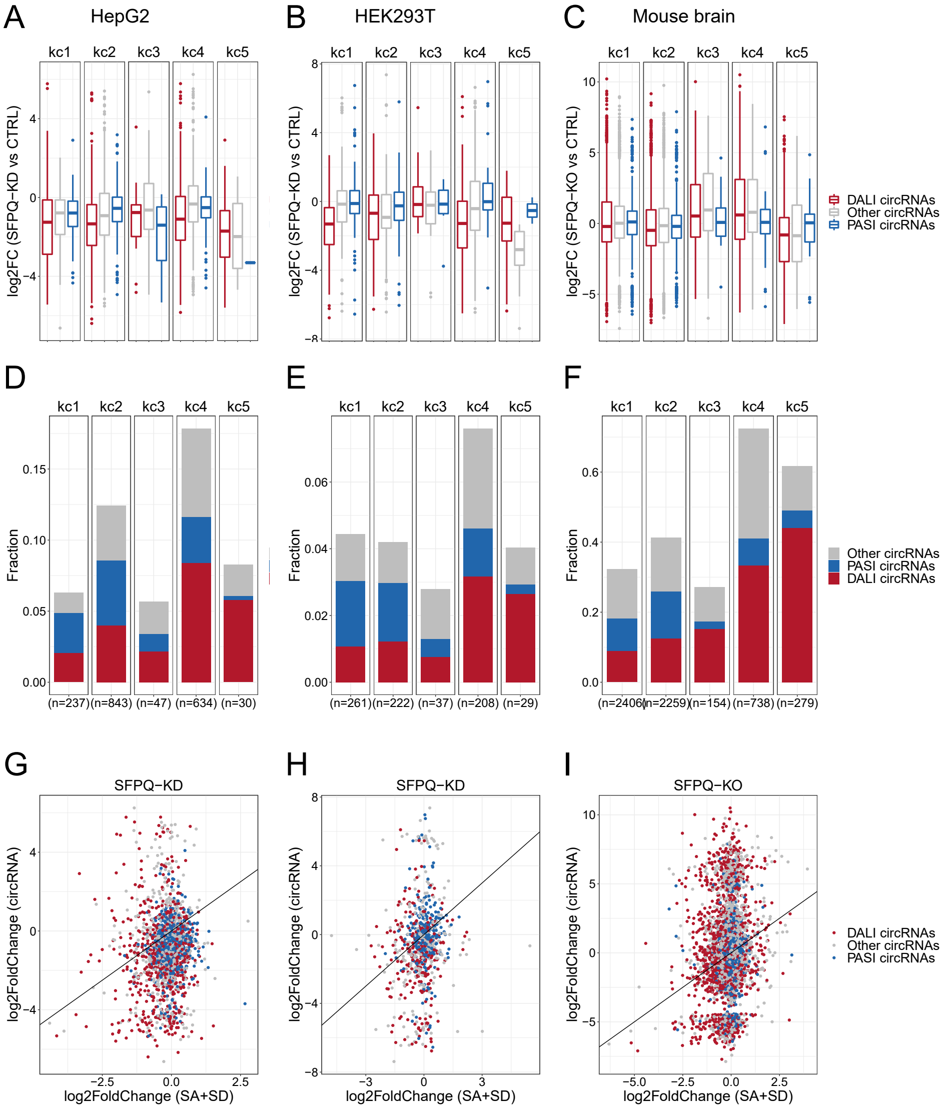


**Supplementary Figure 10: circRNAs in kmeans clusters. A-C)** For each kmean cluster, boxplots showing the log2FoldChange of circRNAs expression upon SFPQ depletion in HEK293T (A), HepG2 (B) cells and mouse brain (C) stratified by circRNAs subgroup. **D-F)** Barplot of numbers and fraction of circRNAs in each kmean cluster in HEK293T (D), HepG2 (E) cells and mouse brain (F). The fraction is determined by the number of genes hosting circRNAs relative to the total number of genes in each cluster. **G-I)** Scatterplot relating the circRNA deregulation (log2FC) with the deregulation of host-gene linear splicing for HEK293T (G), HepG2 (H) cells and mouse brain (I) colorcoded by circRNAs subgroup. The diagonal line represents the perfect correlation.

Supplementary Figure 11

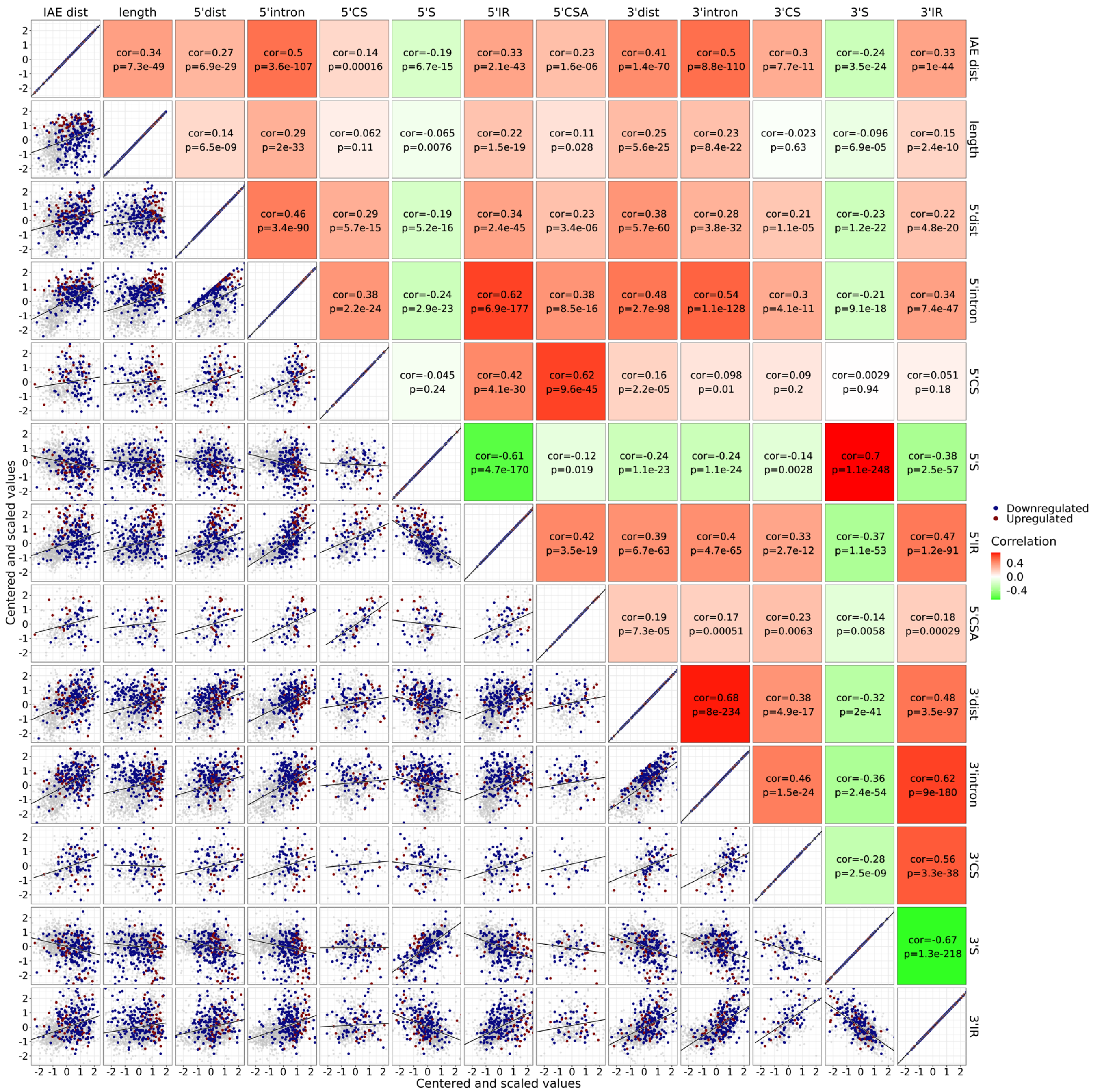


**Supplementary Figure 11: HepG2 features.** Bottom-left; Correlation matrix showing for each pair of features the correlation between standardized (centered and scaled) values. The points are color-coded by circRNA regulations, each significant up (red), significant down (blue) or unchanged (grey). Top-right; the correlation values (based on Pearson correlation) and corresponding p-values are shown. The tiles are color-coded by the correlation values.

Supplementary Figure 12

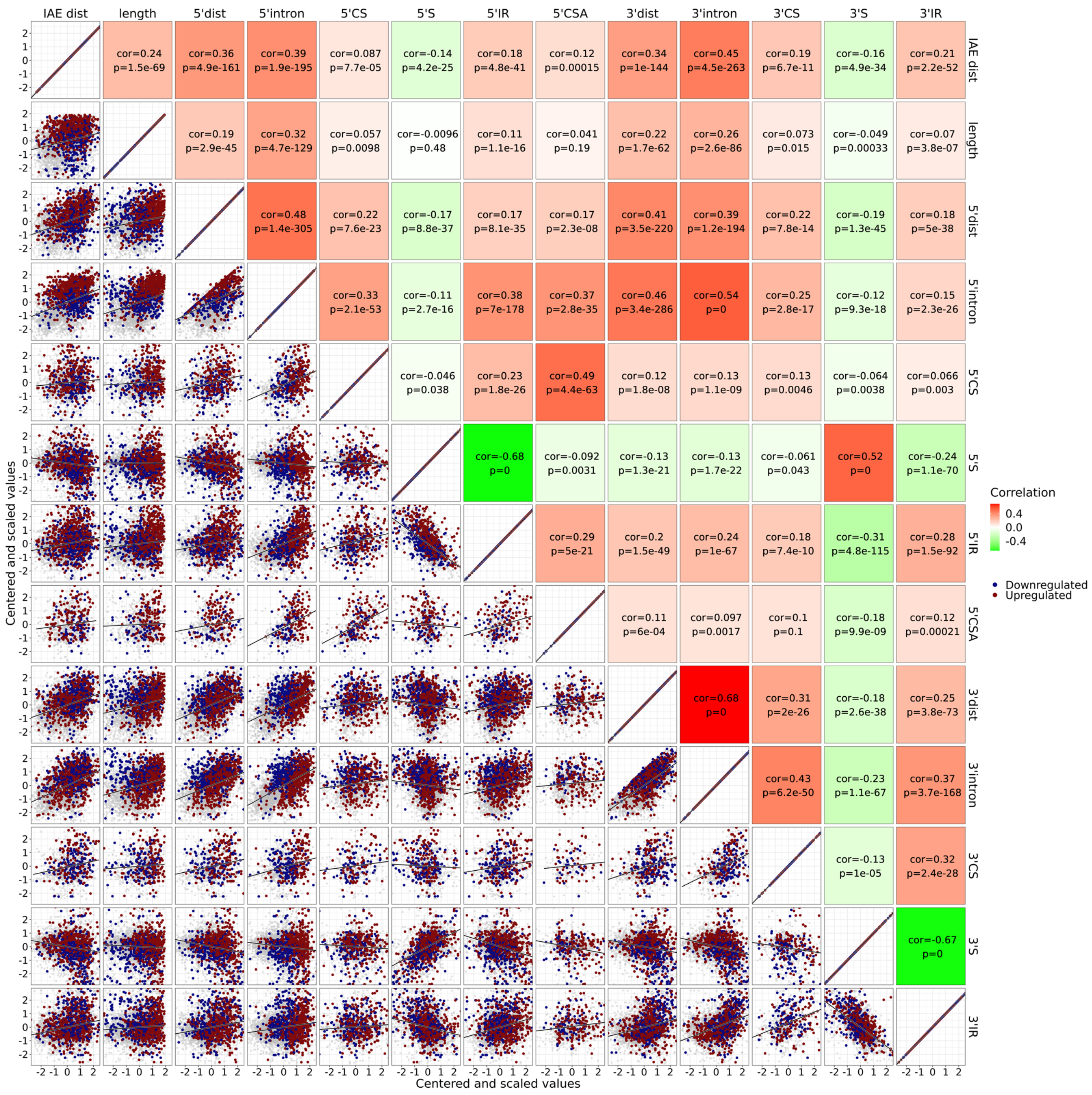


**Supplementary Figure 12: Mouse brain features.** As in Supplementary Fig. 11, but with features from mouse brain data.

Supplementary Figure 13

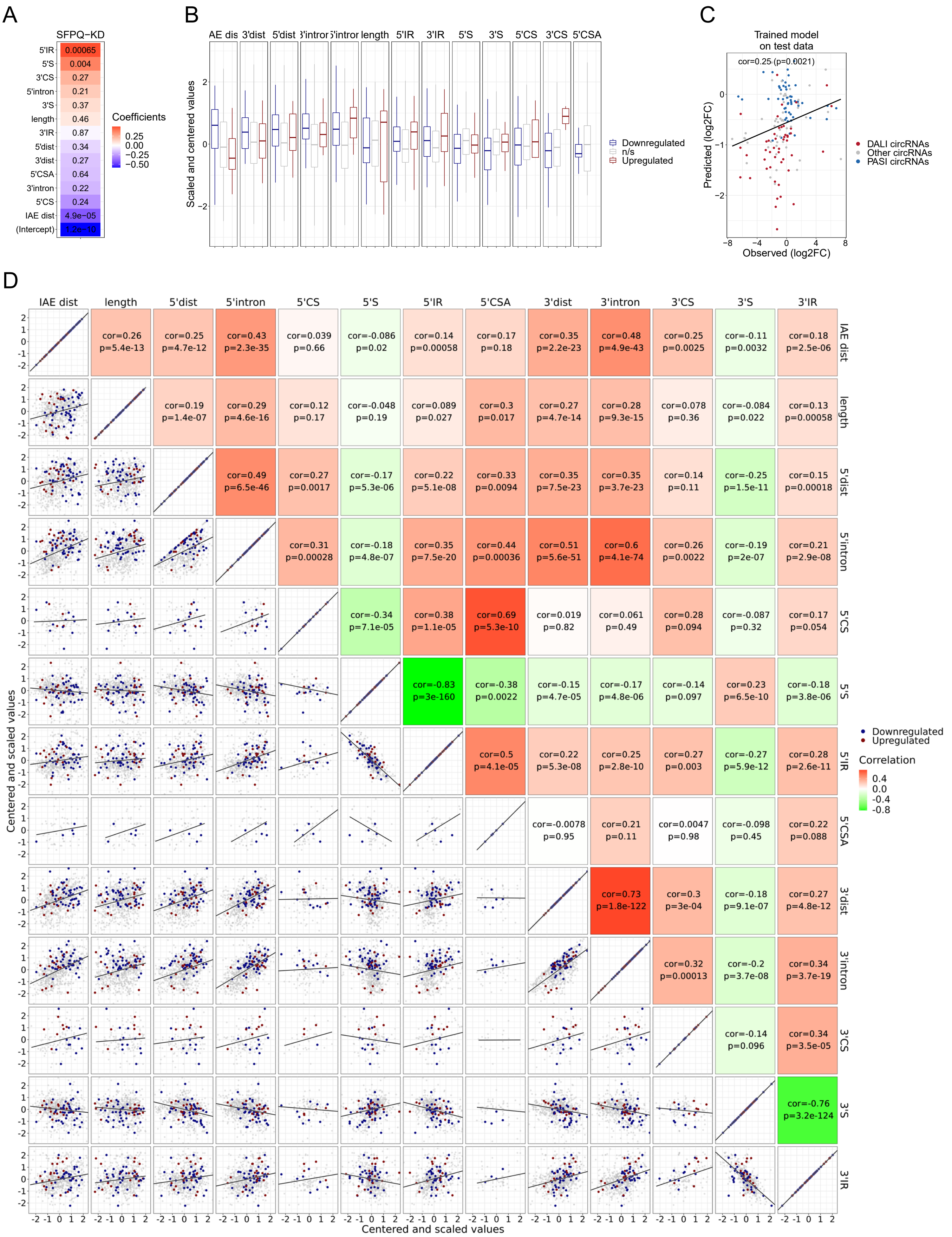


**Supplementary Figure 13: HEK293T features and GLM model performance. A)** As in Figure 6, heat-map showing feature coefficients. **B)** Boxplot showing the standardized feature values for up, down and unchanged circRNAs. **C)** Model prediction on 20% test-set as in Supplementary Fig. 12A. **D)** Correlation matrix as in Supplementary Fig. 11.

Supplementary Figure 14

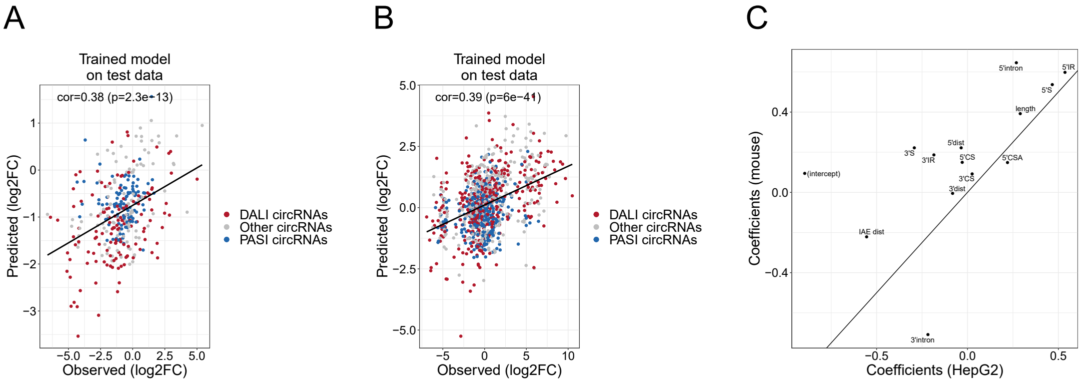


**Supplementary Figure 14: GLM model performance. A-B)** Scatterplot showing the correlation between observed and predicted log2foldchange values on test-set using GLM model in HepG2 (A) and mouse brain (B). The Pearson correlation and corresponding p-value is denoted in the top-left corner. **C)** Scatterplot showing the correlation between GLM coefficients obtain in HepG2 and mouse brain regression analyses.
